# Engineering a Dimeric Single-Domain Antibody for Improved Detection and neutralization of Amyloid-β Oligomers

**DOI:** 10.1101/2025.08.29.672921

**Authors:** Liliana Napolitano, Devkee M. Vadukul, Alessandra Bigi, Cristina Cecchi, Fabrizio Chiti, Roberta Cascella, Francesco A. Aprile

**Affiliations:** Department of Experimental and Clinical Biomedical Sciences, section of Biochemistry, University of Florence, Florence, Italy; Department of Chemistry, Molecular Sciences Research Hub, Imperial College London, London, United Kingdom; Institute of Chemical Biology, Molecular Sciences Research Hub, Imperial College London, London, United Kingdom

**Keywords:** nanobodies, single-domain antibodies, fusion proteins, conformation-sensitive antibodies, amyloid β, oligomers, early-diagnosis, cerebrospinal fluid, neurodegenerative diseases

## Abstract

Soluble Aβ oligomers are regarded as major neurotoxic agents in Alzheimer’s disease. Several monoclonal antibodies have been developed to target Aβ oligomers, but most of them show limited specificity binding also to monomers and fibrils. To generate an antibody with high specificity for the oligomers, we aimed to increase the efficiency and sensitivity of a humanized Aβ-oligomer-specific single domain antibody, called DesAb-O. We engineered a dimeric DesAb-O variant, DiDesAb-O, which showed a significantly improved binding properties for Aβ oligomers as compared to the monomeric sdAb. Furthermore, DiDesAb-O detected Aβ_42_ oligomers in cells, prevented their binding to cell membranes and the Aβ_42_ oligomers-induced neurotoxicity from both synthetic Aβ_42_ samples and cerebrospinal fluid of Alzheimer’s patients at lower concentrations compared to DesAb-O. Overall, our findings indicate that the rational engineering of dimeric sdAb variants is an effective strategy to improve their binding properties offering new opportunities for the development of clinical molecules in the early diagnosis and cure of Alzheimer’s disease.

**Significance Statement:** Protein aggregates, characterized by high structural heterogeneity, are central to numerous neurodegenerative diseases, yet the development of aggregate-specific molecular probes and therapeutic agents remains a significant challenge. We present a novel approach to enhance the binding of single-domain antibodies by engineering multivalent, specifically dimeric, molecules to exploit the avidity effect. As a proof of concept, we developed a dimeric version of a previously studied single-domain antibody named DesAb-O, which exhibits enhanced binding sensitivity and specificity to toxic amyloid-β oligomers and greater efficacy in inhibiting their toxicity. These findings provide a robust foundation for creating next-generation antibody fragments with enhanced binding to heterogeneous protein aggregates, opening new avenues for innovative diagnostic tools and therapeutic strategies in neurodegenerative disease research and treatment.

## Background

Approximately 55 million people worldwide suffer from dementia, with Alzheimer’s disease (AD) being the leading cause accounting for an estimated 60–80% of cases (2024 Alzheimer’s disease facts and figures). This number is predicted to double by 2050 (2024 Alzheimer’s disease facts and figures), thus the need for accurate and early diagnostic tools has become increasingly urgent.

AD is primarily characterized by brain atrophy and synapse loss associated with the accumulation of extracellular amyloid-beta (Aβ) plaques and intracellular tau neurofibrillary tangles (Selkoe and Hardy, 2016; Congdon and Sigurdsson, 2018). Notably, Aβ aggregation begins at least two decades before the appearance of clinical symptoms of AD (Jack et al., 2013; Jansen et al., 2015; Vermunt et al., 2019). Aβ exists in the form of peptides of varying length, ranging from 37 to 49 residues (Nunan and Small, 2000). The deposition of amyloid fibrils of the 42-residue Aβ (Aβ_42_) in specific brain areas as neuritic plaques represents a major pathological hallmark of AD (Bloom, 2014), and an early histopathological feature revealed by the earliest biomarkers, such as amyloid plaques with by position emission tomography (PET) or Aβ_42_ depletion from the cerebrospinal fluid (CSF) (Jack et al. 2013). These features precede the clinical diagnosis of dementia by ca. 5-15 years (Vermunt et al., 2019).

While Aβ_42_ fibrils are a pathological feature of AD, growing evidence suggests that it is the smaller, soluble aggregation intermediates, *i.e.*, the oligomers, that are primarily responsible for the neurotoxic effects observed in AD (Kayed et al., 2003; Benilova et al., 2012; Selkoe and Hardy, 2016; Selkoe, 2019; Gutierrez and Limon, 2022). Increased oligomer levels have been detected in the brain and CSF of AD patients compared to age-matched controls (Kayed et al., 2003; Gong et al., 2003; Lasagna-Reeves et al. 2011; Hefti et al., 2013; Savage et al., 2014; Tooyama et al., 2023) and have shown to correlate stronger with the severity of cognitive decline and with the loss of synaptic markers compared to mature amyloid fibrils (Cline et al., 2018; Selkoe, 2019). Despite the prevalence of oligomers in AD, suitable sensitive methods for their routine and reliable detection, quantification and isolation from biological fluids have been lacking due to their transient and heterogeneous nature.

Recent progress in the development of conformation-specific monoclonal antibodies (mAbs) has enabled the targeting of toxic Aβ_42_ oligomers (Kayed et al. 2003; Gong et al. 2003; Lasagna-Reeves et al. 2011; Savage et al., 2014; Murakami et al., 2014; Tooyama et al., 2023). Additionally, mAbs have been developed to target Aβ in AD patients as a therapeutic strategy (Karran and De Strooper, 2022), with several currently undergoing clinical trials (Sevigny et al., 2016; Swanson et al., 2021). Among them, Aducanumab (Budd Haeberlein et al. 2022; Chen et al., 2024), Lecanemab (van Dyck et al., 2023; Swanson et al., 2021) and Donanemab (Sims et al., 2023) have received FDA approval for AD treatment, although production of the former has been later discontinued by its producer company Biogen. Use of these mAbs in the clinical practice has been associated with adverse effects, including intracerebral hemorrhages and Amyloid-Related Imaging Abnormalities (ARIA) (van Dyck et al., 2023; Sims et al. 2023; Cummings et al., 2024).

The discovery of conformation-sensitive antibodies (Abs) and the approval of monoclonal antibodies (mAbs) as the first disease-modifying therapy for AD have been significant breakthroughs. However, their severe side effects represent a significant limitation for clinical application. On the contrary, single-domain Abs (sdAbs), formed only by the variable fragment of the heavy chain of camelid antibodies, are emerging as promising candidates for AD diagnosis and therapy, due to their small size, high solubility and low immunogenicity (Lafaye et al., 2009; Muyldermans, 2013; Ackaert et al., 2021; De Leiris et al., 2024). Previously published work demonstrated that targeting the region 29 to 36 of Aβ_42_ with a human rationally designed sdAbs on the VH domain of the clinically approved mAb Trastuzumab (Baselga et al., 1998; Barthelemy et al., 2008), called DesAbs, can interfere with the Aβ aggregation process (Aprile et al., 2017). Among them, DesAb-O was found to preferentially bind to Aβ oligomers rather than the monomers or fibrils, as this region is likely to be solvent-exposed when the peptide is oligomeric, before becoming buried in mature fibrils (Aprile et al., 2020). In a further study, DesAb-O was found to be able to selectively detect synthetic Aβ_42_ oligomers both *in vitro* and in cultured cells, and to prevent their associated neuronal dysfunction (Bigi, Napolitano et al., 2024). Notably, DesAb-O could also identify and neutralize Aβ_42_ oligomers present in the CSF samples (CSFs) of AD patients with respect to healthy individuals (Bigi, Napolitano et al., 2024).

Recent studies have shown that multivalency is an effective strategy to enhance the effects of sdAbs, leading to improved protein detection, increased avidity and binding ability for protein aggregates (Zhao et al., 2023; McArthur et al., 2024). Building on this concept, we engineered a dimeric variant of DesAb-O, named DiDesAb-O, to further enhance its binding properties and refine its ability to detect and neutralize oligomers. Our data show that DiDesAb-O has an enhanced anti-aggregation activity and sensitivity to detect Aβ_42_ oligomers *in vitro* and in cultured cells as compared to the monomeric counterpart DesAb-O, and is able to neutralize toxic species in both synthetic Aβ_42_ samples and AD CSF at lower concentrations as compared to DesAb-O. Our findings also demonstrate that the rational engineering of dimeric sdAbs is a promising avenue for developing potential diagnostic and therapeutic molecules targeting AD and other neurodegenerative diseases.

## Materials and methods

### DiDesAb-O design

To design DiDesAb-O, a flexible linker composed by a repetition of glycine (Gly) and Serine (Ser), was attached to the C-terminus of a first DesAb-O monomer and the N-terminus of a second DesAb-O monomer, depleted of the initial four amino acids (Methionine (Met), Arginine (Arg), Gly and Ser), and the following 6X His-tag region. The precise sequence of the linker was GGGGSGGGGSGGGGS. Gene sequence design was carried out through the use of SnapGene software (www.snapgene.com).

### Preparation of amyloid β-derived diffusible ligands (ADDLs)

The lyophilized synthetic Aβ_42_ peptide (Bachem) was dissolved in 100% hexafluoro-2-isopropanol (HFIP) to obtain a monomeric form, followed by evaporation under nitrogen flux. To obtain ADDLs, Aβ_42_ was resuspended in anhydrous dimethyl sulfoxide (DMSO) to 5 mM and then diluting in ice-cold F-12 medium to a final concentration of 100 μM. This solution was incubated at 4 °C for 1 day and then centrifuged at 14,000 × g for 10 min, as previously reported (Lambert et al., 1998).

### Thioflavin T Fluorescence Assays

Monomeric recombinant Aβ_42_ (1 μM) was incubated in the absence or presence of DiDesAb-O at decreasing molar ratios (1:1, 1:0.5, 1:0.25, 1:0.125, corresponding to 1 μM, 0.5 μM, 0.25 μM, 0.125 μM DiDesAb-O concentrations) in 20 mM phosphate buffer, pH 8.0, 37 °C. Samples were prepared with a final concentration of 10 μM Thioflavin T (ThT) dye, gently vortexed, and pipetted into nonbinding surface black 96-well plates (Greiner Bio-One) in triplicates. The plate was read in a ClarioStar Plus microplate reader (BMG LabTech) at 37 °C. The excitation and emission wavelengths were set to 440 and 480 nm, respectively, and fluorescence intensity measurements were taken using spiral averaging (3 mm diameter). Buffer-only values were not subtracted from the sample readings but shown in the raw data analysis. Readings were taken every 2 min. To test the capability of DiDesAb-O compared to DesAb-O, in another set of experiments monomeric Aβ_42_ (1 μM) was incubated alone or in presence of 1 μM DiDesAb-O or increasing DesAb-O molar ratios (1:1, 1:2 corresponding to 1 and 2 μM DesAb-O concentrations) under the same conditions. The data were plotted using GraphPad Prism version 9.3.1 for Windows (GraphPad Software).

### Real-Time ELISA

Real-time ELISA experiments were performed aggregating 1 μM monomeric recombinant Aβ_42_ peptides in 20 mM sodium phosphate buffer, pH 8.0 in quiescent conditions, at 37 °C. Aliquots of 20 µl were taken at precise timepoints (0, 0.5, 1, 2 and 20 h) from aggregation reactions and immobilized on a 96- or 384-well Maxisorp ELISA plate (Nunc) with no shaking overnight at 4 °C. Aβ_42_ fibrils obtained after 4 days of incubation at 37 °C were used as a control. At the end of the incubation, the plate was then washed three times with TBS (20 mM Tris, pH 7.4, and 100 mM NaCl) and incubated in TBS supplemented with 5% bovine serum albumin (BSA) under constant shaking for 1 h at RT. The plate was then washed six times with TBS and then incubated with 30 μL solutions 1 μM DesAb-O or 1 μM DiDesAb-O under constant shaking either for 1 h at RT or overnight at 4 °C. At the end of this incubation, additional six washes with TBS were performed and the plate was incubated with 30 μL solutions of rabbit polyclonal 6X His tag horseradish peroxidase (HRP) conjugated (Abcam) at a dilution of 1:4.000 in 20 mM Tris, pH 7.4, 100 mM NaCl, and 5% BSA under constant shaking for 1 h at RT. Finally, the plate was washed twice with 20 mM Tris, pH 7.4, and 100 mM NaCl, then three times with 20 mM Tris, pH 7.4, 100 mM NaCl, and 0.02% Tween-20 and again three times with 20 mM Tris, pH 7.4, and 100 mM NaCl. Finally, the amount of bound sdAbs was quantified by using 1-Step Ultra TMB-ELISA Substrate Solution (Thermo Fisher Scientific), according to the manufacturer’s instructions, and the absorbance was measured at 450 nm by means of a CLARIOstar plate reader (BMG Labtech) as previously reported (Aprile et al., 2020).

### Transmission Electron Microscopy (TEM)

Aβ_42_ recombinant samples incubated at 5 μM in the absence or in the presence of equimolar concentration of sdAbs were induced to aggregate in 20 mM sodium phosphate buffer, pH 8.0, in a microplate without ThT and collected after 24 h of aggregation at 37 °C. Aβ_42_ aggregation was monitored by other replicates with ThT, allowing real-time monitoring of the reaction. Samples for transmission electron microscopy (TEM) were then prepared spotting 4 μL onto Formvar/carbon-coated 300 mesh copper grids for 1 min. By blot drying the grid with Whatman filter, we removed excess sample, allowing the grid drying for 2 min. Samples were then washed with 4 μL of water and stained with 4 μL of 2% w/v uranyl acetate (Vadukul et al., 2023) and the grid was dried again as described above. Grids were imaged on a T12 Spirit electron microscope (Thermo Fisher Scientific). The diameters of various fibrils were measured using ImageJ software and all data were plotted using Excel (Version 16.89.1 (24091630)).

### Dot Blot

Dot blots were carried out on 5 μM recombinant Aβ_42_ samples that were aggregated in 20 mM phosphate buffer, pH 8.0, at 37 °C in the microplates without ThT for 4 d. Samples were collected and centrifuged at max speed (∼17,000g) for 1 h on a benchtop centrifuge to separate the soluble (supernatant) and insoluble aggregates (pellet). 8 μl of each sample were collected before centrifugation as total protein quantity. At the end of the centrifuge, the supernatant was collected and the pellet was resuspended in 20 mM phosphate buffer, pH 8.0. Three repeats of each sample were spotted onto a 0.45 μM nitrocellulose membrane and blocked in 5% non-fat milk in 0.1% PBS-Tween for 1 h at RT. The membranes were then incubated with 1:1000 diluted mouse monoclonal anti-Aβ (6E10) Ab (Biolegend) or 1:1000 diluted anti-6X His-tag Ab (Abcam) in 0.1% PBS-Tween overnight at 4 °C, under constant shaking. The following day, the membranes were washed three times for 10 min each with 0.1% PBS-Tween and incubated with anti-mouse Alexa Fluor (AF) 647-conjugated in 0.1% PBS-Tween for 1 h at RT, protected against light. Following three further washes for 10 min each in 0.1% PBS-Tween, the membranes were detected with the appropriate laser using a Typhoon scanner (GE Healthcare).

### CSF samples

CSF samples (CSFs) from human aged controls (n = 4) or AD patients (n = 4) were obtained from BioIVT. Each CSF was received in 0.5–1 ml aliquot and stored at – 80 °C. Samples were centrifuged at 4000 g for 10 min at 4 °C, obtaining a pale pellet that was separated from the supernatant as previously reported (Bigi, Napolitano et al., 2024). The supernatant was then analysed; protein concentration in these samples was determined by the Bradford colorimetric method (Bradford MM, 1976).

### Cell cultures

Authenticated human SH-SY5Y neuroblastoma cells were purchased from A.T.C.C. and cultured in Dulbecco’s modified Eagle’s medium (DMEM), F-12 Ham with 25 mM 4-(2- Hydroxyethyl) piperazine-1-ethanesulfonic acid (HEPES) and NaHCO_3_ (1:1) supplemented with 10% fetal bovine serum (FBS), 1.0 mM glutamine and 1.0% penicillin and streptomycin solution (Sigma-Aldrich), as reported previously (Capitini et al. 2014; Bigi, Fani et al., 2024). Cells were maintained in a 5.0% CO_2_ humidified atmosphere at 37 °C and grown until 80% confluence for a maximum of 20 passages and tested to ensure that they were free form mycoplasma contamination (Bigi, Fani et al., 2024; Bigi, Napolitano et al., 2024).

### Confocal microscopy

Synthetic Aβ_42_ oligomers obtained after 8 h of incubation at a concentration of 10 µM in PBS, pH 7.4, 37 °C, were added to the culture medium of SH-SY5Y cells seeded on glass coverslips for 1 h at 0.5 µM. After incubation, the cells were washed with PBS, the plasma membranes were counterstained with 0.01 mg/ml tetramethylrhodamine-conjugated wheat germ agglutinin (WGA; Thermo Fisher Scientific) for 15 min at 37 °C and cells were then fixed with 2.0% (w/v) paraformaldehyde for 10 min at RT, as reported (Cascella et al., 2021; Bigi, Napolitano et al., 2024). After washing with PBS, the plasma membranes were permeabilized with a 3.0% (v/v) glycerol solution for 10 min at RT. Aβ_42_ species were then detected with 3 μM and 1 μM DesAb-O or DiDesAb-O and 1:800 6E10 Ab as a control. As a secondary Ab, we used diluted FITC anti-6X His-tag secondary Abs (Abcam) or AF488-conjugated anti-mouse secondary Abs (Thermo Fisher Scientific).

In another set of experiments, we calculate the number of Aβ_42_ oligomers bound to neuronal membranes of SH-SY5Y. To perform this experiment, 0.5 μM Aβ_42_ oligomers were pre-incubated or not with decreasing sdAbs molar ratios (1:3, 1:2, 1:1, 1:0.5, 1:0.25 and 1:0.1 corresponding to 7.5, 5, 2.5, 1.25, 0.63 and 0.25 µM of sdAbs concentrations) for 1 h at 37 °C under soft shaking, and cells were then treated for 15 min. At the end of treatment, cells were washed with PBS and the plasma membranes were counterstained for 15 min with 5.0 μg mL−1 AF633-conjugated WGA (Thermo Fisher Scientific) (Bigi, Napolitano et al., 2024). Cells were fixed in 2.0% (w/v) paraformaldehyde for 10 min at RT and Aβ_1–42_ assemblies were detected with 1:800 diluted 6E10 Ab and then 1:1000 diluted AF488-conjugated anti-mouse secondary Ab (Thermo Fisher Scientific). To detect only the oligomers bound to the cell surface, the cellular membrane was not permeabilized, thus preventing Ab internalization. Fluorescence emission was detected after double excitation at 633 and 488 nm by a TCS SP8 scanning confocal microscopy system (Leica Microsystems), as previously described (Bigi et al., 2020). The number of oligomers was estimated in regions of interest in 60-80 cells, via the use of ImageJ software (NIH) and JACOP plugin (http:// rsb. info. nih. gov) (Rasband WR).

### Measurement of cytosolic free Ca^2+^ levels

The intracellular calcium levels were measured in SH- SY5Y cells as previously described (Bigi et al., 2023, Bigi, Napolitano et al., 2024). SH-SY5Y cells were treated for 15 min with 0.5 μM synthetic Aβ_42_ oligomers previously incubated or not for 1 h at 37 °C under gentle shaking with decreasing sdAbs molar ratios (1:3, 1:2, 1:1, 1:0.5, 1:0.25 and 1:0.1 corresponding to 7.5, 5, 2.5, 1.25, 0.63 and 0.25 µM sdAb). Cells were then washed in PBS and loaded with 4.5 μM Fluo-4 AM (Thermo Fisher Scientific) for 10 min and cytosolic Ca^2+^ levels were detected after excitation at 488 nm by the TCS SP8 scanning confocal microscopy system as previously reported (Cascella et al., 2021; Bigi et al., 2023; Bigi, Napolitano et al., 2024).

In another set of experiments, SH-SY5Y cells were treated for 5 h with the CSFs (n = 4 for AD as well as for controls) pre-incubated or not with DesAb-O or DiDesAb-O at 3 μM and 1 μM for 1 h at 37 °C under shaking, as previously described (Bigi, Napolitano et al., 2024). Images were then analyzed using the ImageJ software, and the fluorescence intensities were expressed as the percentage of that measured in untreated cells, taken as 100%.

### MTT reduction assay

To assess the cytotoxicity of Aβ_42_ oligomers, a 3-(4,5-dimethyltohiazol-2- yl)-2,5-diphenyltetrazolium bromide (MTT) reduction assay was conducted. Briefly, 8 h Aβ_42_ oligomers were pre-incubated for 1 h at 37 °C with decreasing sdAbs molar ratios (1:3, 1:2, 1:1, 1:0.5, 1:0.25 and 1:0.1 corresponding to 7.5, 5, 2.5, 1.25, 0.63 and 0.25 µM sdAbs) and then added to the extracellular medium of SH-SY5Y cells. After treatment, the cell culture medium was removed, cells were washed in PBS and the MTT solution was added to the cells for 4 h. The formazan product was solubilized with cell lysis buffer (20% sodium dodecyl sulphate (SDS), 50% N, N-dimethylformamide, pH 4.7) for 1 h. The absorbance values of blue formazan were determined at 595 nm. MTT tests were achieved using Microplate Manager® Software (Biorad). Cell viability was expressed as the percentage of MTT reduction relative to untreated cells (taken as 100%), or to those treated with Aβ_42_ species in the absence of sdAbs, as previously reported (Bigi, Napolitano et al., 2024).

### Statistical analysis

Data were expressed as means ± standard deviation (S.D.), or as means ± standard error of mean (S.E.M), as indicated in each figure legend. Comparisons between the different groups were performed by using one-tailed *Student* t-test or by One-way or Two-way ANOVA followed by Bonferroni’s multiple-comparison test (GraphPad Prism 10.3.1 software), as indicated in each figure legend. P values lower than 0.05, 0.01, 0.001 and 0.0001 were considered to be statistically significant (*), highly statistically significant (**), very highly statistically significant (***), and extremely highly statistically significant (****), respectively.

## Results

### DiDesAb-O design strategy

The rationale behind the engineering of DiDesAb-O was to enhance the binding to the Aβ_42_ oligomers by increasing the avidity of the monomeric DesAb-O previously studied (Aprile et al., 2020; Bigi, Napolitano et al., 2024). It is conceivable that the dimer may be capable of binding to multiple exposed binding sites within the oligomeric structure. This could result in an enhanced binding avidity for Aβ_42_ oligomers. To obtain DiDesAb-O, the C-terminus of one DesAb-O monomer was linked to the N-terminus of another DesAb-O monomer *via* a flexible linker, consisting of three repeats of the Gly-Gly-Gly-Gly-Ser sequence (Huston et al., 1988), as illustrated in **Figure 1A**. Notably, DesAb-O N-terminus contains residues that are preferred in protein linker regions (Argos, 1990; Chen et al., 2013), allowing it to function as an extension of the linker itself and effectively increasing the overall linker length (**Figure 1A**). This extended linker enhances the flexibility and ability of DiDesAb-O to bind distant epitopes, such as those found in the Aβ_42_ oligomers (Chen et al., 2013; Yusakul et al., 2016). The entire amino acid sequence of DiDesAb-O is shown in **Figure S1**.

**Figure 1.**
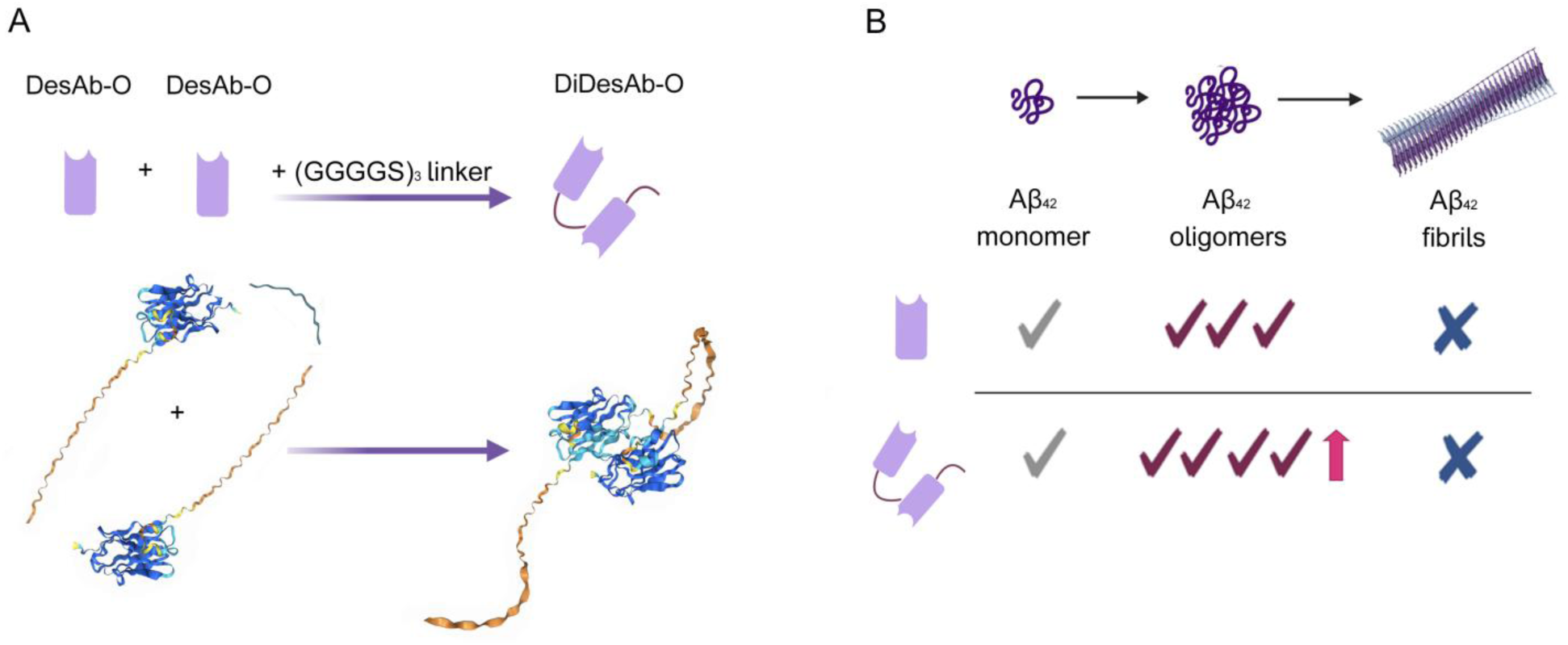
DiDesAb-O design. **A** Schematic representation of the procedure to obtain the DiDesAb-O (top). On the left, two monomers of DesAb-O and the (GGGGS)_3_ flexible linker (blue) are shown. On the right, the DiDesAb-O structure is represented as the results of the union of the previous three components. DesAb-O, DiDesAb-O and (GGGGS)_3_ flexible linker reconstructions were obtained with AlphaFold 3. **B** Schematic representation of the aim of the study. We designed a dimeric structure of DesAb-O to obtain an enhanced sensitivity and binding avidity for Aβ_42_ oligomers, while avoiding increased binding to Aβ_42_ monomers or fibrils.

### Structural characterization of DiDesAb-O

DiDesAb-O was expressed in *E. coli* and purified, as detailed in the *Materials and Methods* section and in **Figure S2**. The molecular weight of the purified construct, its secondary structure and thermal stability were characterized by far-UV circular dichroism (CD) and electrospray ionisation mass spectrometry (ESI-MS), as reported in **Figure S3**. Far-UV CD spectroscopy showed that DiDesAb-O exhibits a predominant β-sheet secondary structure, similar to that observed for the monomer (**Figure S3A**).

To provide a quantitative interpretation of the far-UV CD spectrum of DiDesAb-O and compare it with that previously obtained for DesAb-O (Aprile et al., 2020), the BestSel software (v1.3.230210; Micsonai et al., 2018) was employed (**Table S1**). BestSel analysis yielded similar proportions of antiparallel β-sheet secondary structure for DiDesAb-O compared to DesAb-O (61% and 57%, respectively). Notably, DesAb-O was found to comprise 7% parallel β-sheet, an element that is entirely absent from the dimeric structure profile. Overall, the total β-sheet structure is similar for dimeric and monomeric sdAbs (61% and 64%, respectively). Conversely, the dimeric sdAb was found to exhibit a 3% α-helix, possibly attributable to the presence of the linker, which is absent for DesAb-O. The quantification of turns plus disordered secondary structure is similar in the two cases, amounting to 36%. Based on the ESI-MS spectrum, the mass of the purified protein was determined to be 34,270 Da (**Figure S3B**). This experimental value closely matched the expected mass of DiDesAb-O, calculated as 34,269 Da using the ExPASy ProtParam tool (Gill and Von Hippel, 1989).

To assess whether DiDesAb-O had similar structural stability to DesAb-O, we performed a thermal denaturation experiment to determine the temperature of half-denaturation (*T_m_*) (**Figure S3C,D**). CD spectra were recorded between 20 °C and 90 °C. The mean molar residue ellipticity per residue ([θ]_res_) at 210 nm was extracted from each CD spectrum and plotted as a function of temperature. The data were converted into fraction folded (%) values and fitted with a two-state model. The two sdAbs fragments had similar *T_m_* values (73.7 °C for DiDesAb-O and 74.6 °C for DesAb-O). The slight decrease of *T_m_*value and cooperativity for DiDesAb-O relative to DesAb-O likely results from the high net charge of DesAb-O monomers that generates a further repulsion between the two monomeric units in DiDesAb-O, with consequent destabilisation (lower *T_m_* value) and lower enthalpy change at the *T_m_*(*ΔH_m_*) resulting in lower transition cooperativity.

This structural characterization showed how the engineering of DesAb-O and the addition of a flexible linker resulted in a stable dimeric sdAb that shared a secondary structure and thermal stability close to those of the corresponding monomeric sdAb.

### DiDesAb-O slows down Aβ_42_ aggregation more effectively than DesAb-O

To assess the ability of DiDesAb-O to interfere with Aβ_42_ aggregation, we performed ThT assays on solutions containing 1 μM monomeric Aβ_42_ in the presence of various Aβ_42_:DiDesAb-O molar ratios (1:1, 1:0.5, 1:0.25, 1:0.125), as reported in **Figures 2A and S4A**. Solutions containing only the highest DiDesAb-O concentration (1 μM) in the absence of Aβ_42_ were tested as a control (**Figure S4A**). DiDesAb-O significantly reduced the Aβ_42_ aggregation process at the 1:1 molar ratio, increasing the aggregation half-times (*t_50_*) by approximately 2-fold compared to Aβ_42_ alone (3.2 ± 1.1 h and 1.5 ± 0.4 h, respectively). Furthermore, the *t_50_* value decreased as a function of DiDesAb-O concentration (**Table S2**) demonstrating the ability of DiDesAb-O to interfere with the Aβ_42_ aggregation even at sub-stochiometric concentrations (**Figure 2A**). *t_50_* values were determined as the mean of three independent experiments (**Table S2**), whereas **Figures 2A and S4A** only show one representative kinetic trace per condition.

**Figure 2.**
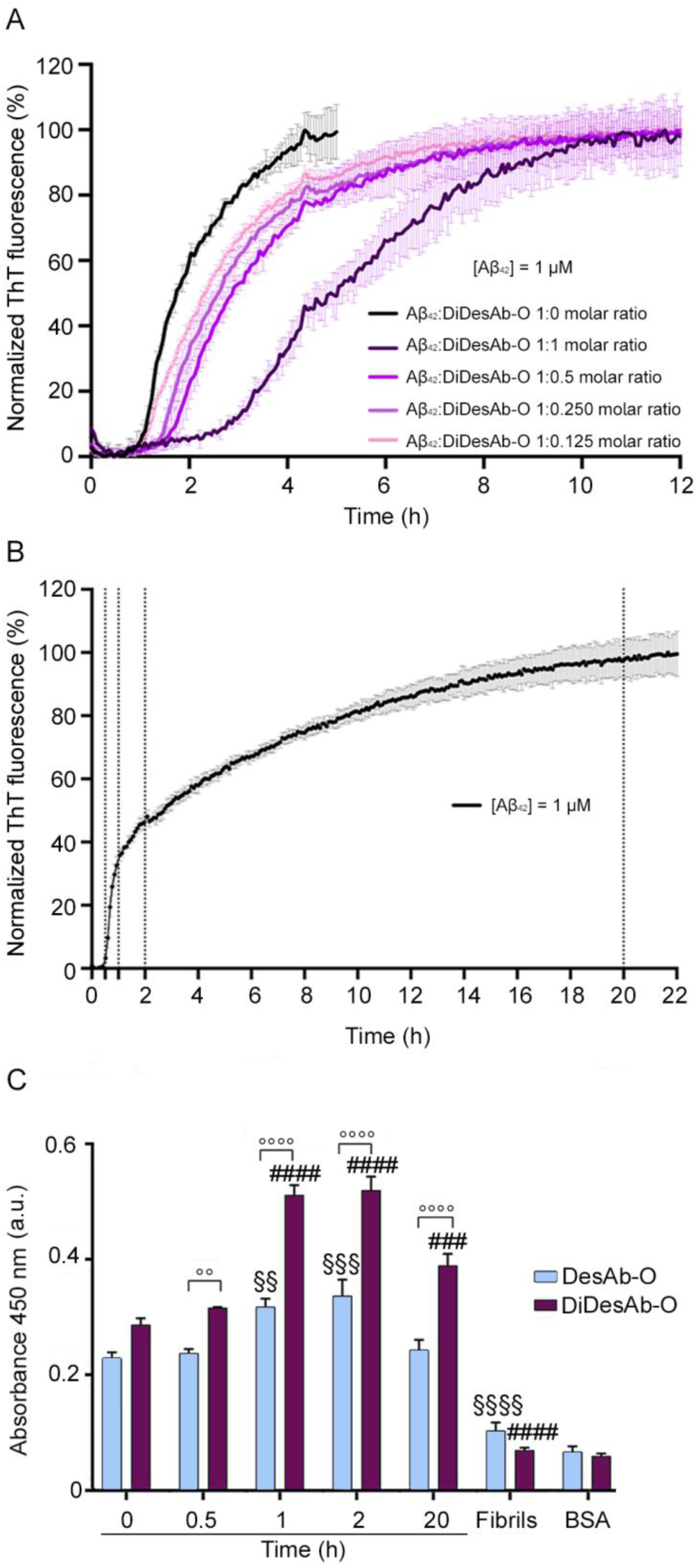
DiDesAb-O interferes with the Aβ_42_ aggregation process and has improved binding properties for Aβ_42_ oligomers. **A** Representative ThT fluorescence assays of 1 μM Aβ_42_ at the indicated Aβ_42_:DiDesAb-O molar ratios: 1:0 (black), 1:1 (dark purple), 1:0.5 (bright purple), 1:0.25 (dark pink) and 1:0.125 (pink). **B** ThT-based *in vitro* aggregation assay of 1 μM Aβ_42_ in the absence of DiDesAb-O (average of three replicates is shown). The black dashed lines indicate the timepoints at which samples were collected from the aggregation reaction to perform the ELISA experiment. In both panels, error bars refer to standard deviations. **C** ELISA experiment performed on samples collected from the aggregation reaction shown in **B**, using DesAb-O and DiDesAb-O as primary Abs. Aβ_42_ fibrils obtained after 4 days of incubation at 37 °C were used as a control. BSA signals represent the background absorbance values. The bar corresponding to DesAb-O is coloured light blue while the one corresponding to DiDesAb-O is purple. Error bars indicate S.E.M. Statistical analysis was performed by ANOVA with multiple comparison. Samples (n = 4) were analysed by Two-way ANOVA multiple comparison test comparing the absorbance obtained with DesAb-O and DiDesAb-O at the same timepoints (°°P <0.01 and °°°°P<0.0001) and versus their respective values at time 0 min (§§P <0.01, §§§P <0.001 and §§§§P<0.0001 for DesAb-O and ###P < 0.001 ####P < 0.0001 for DiDesAb-O).

To evaluate the effect of DiDesAb-O with respect to DesAb-O, we tested increasing molar ratios of Aβ_42_:DesAb-O (1:1, 1:2) and compared the kinetic traces with those obtained with DiDesAb-O, as represented in **Figure S4A,B**. The aggregation profiles obtained with increasing DesAb-O molar ratios slightly increased the *t_50_* value (1.8 ± 0.6 h for 1:2, 1.7 ± 0.4 h for 1:1, 1.5 ± 0.4 h for 1:0 Aβ_42_:DesAb-O, respectively), slowing down the Aβ_42_ aggregation process (**Figure S4B,C**). Despite this interference, DesAb-O did not slow the aggregation process to the same extent as DiDesAb-O, increasing the *t_50_* value only by 1.3-fold at 1:2 Aβ_42_ monomer:sdAb molar ratio (**Figure S4A,B,C**). Notably, DiDesAb-O exhibited a comparable inhibitory effect at a significantly lower molar ratio of 1:0.25 (*t_50_*value of 1.8 ± 0.5 h), indicating a substantially greater potency of DiDesAb-O in interfering with the aggregation process of Aβ_42_. Again, all *t_50_* values were determined as the mean of three independent experiments, wherein **Figures S4B,C** only shows one representative trace per condition.

### DiDesAb-O has enhanced binding sensitivity for Aβ_42_ oligomers relative to DesAb-O

To compare the binding properties of DiDesAb-O and DesAb-O for Aβ_42_ oligomers, we performed a real-time ELISA. To do so, we prepared solutions of 1 μM Aβ_42_, and we monitored the aggregation of Aβ_42_ by ThT fluorescence (**Figure 2B**). At different timepoints (0, 0.5, 1, 2, 22 h) samples were collected and adsorbed onto an ELISA plate *overnight*. The following day, 1 μM DesAb-O or 1 μM DiDesAb-O were used as primary Abs. Immediately after starting the aggregation reaction (0 h), the absorbance was already higher than that found for BSA used as a background signal, probably due to the detection of low molecular weight Aβ_42_ aggregate and, although to a lesser extent, Aβ_42_ monomer by sdAbs (**Figure 2C**). After 0.5 h, DiDesAb-O slightly increased the absorbance signal probably due to the increased formation of low molecular weight aggregates, exhibiting a statistically difference with DesAb-O at the same time (**Figure 2C**). As the time-course continues, our results revealed a significant increase in absorbance signal for both sdAbs, especially after 1 and 2 h from the beginning of the aggregation reaction, approximately close to the half-time of aggregation as determined by ThT fluorescence (**Figure 2B,C**), where oligomers are at their maximum concentration, due to their role as crucial intermediates in fibril formation (Dobson, 1999; Chia et al., 2018) At these timepoints, DiDesAb-O exhibited the highest absorbance, being approximately 2-fold higher than that of the initial aggregation timepoint, showing a significant difference compared to DesAb-O at the same timepoints. After 20 hours of aggregation, DiDesAb- O still showed an absorbance approximately 1.3-fold higher than that of time 0, indicating the binding to Aβ_42_ oligomeric species, while DesAb-O returned to similar absorbance levels as the initial ones, as previously reported (Aprile et al., 2020).

In order to understand whether the dimeric form of DesAb-O was recognising Aβ_42_ oligomers or Aβ_42_ fibrillar conformers, we used Aβ_42_ fibrils obtained after 4 days of incubation at 37 °C as a control, as represented in **Figure 2C**. Interestingly, neither sdAbs recognised these Aβ_42_ fibrils, resulting in an absorbance signals well below the corresponding values at 0 h and similar to those obtained with BSA. From this evidence, we can assess that DiDesAb-O is able to recognise Aβ_42_ oligomers at very low concentrations, with high specificity and increased sensitivity, representing a successful improvement of the previous DesAb-O binding properties for Aβ_42_ oligomers.

### DiDesAb-O induces morphological changes and increased structural weakness in Aβ_42_ fibrils

To further investigate the impact of DiDesAb-O on Aβ_42_ aggregation, we analyzed the morphological changes and fragility alterations induced by the sdAb in Aβ_42_ fibrils. To do so, we aggregated samples containing 5 μM Aβ_42_ in the absence or presence of an equimolar concentration of either DesAb-O or DiDesAb-O for 4 days at 37 °C. Then, we visualized the samples by transmission electron microscopy (TEM) (**Figure 3A**). Aβ_42_ aggregated in the absence of sdAbs forming a dense network of long fibrils with a diameter of 11.5 ± 0.1 nm (**Figure 3A,B)**. In contrast, Aβ_42_ fibrils obtained in the presence of DesAb-O were shorter and exhibited a reduced dimeter of 9.1 ± 0.8 nm (**Figure 3A,B**). A striking change in the morphology of Aβ_42_ fibrils was observed when monomeric Aβ_42_ was co-incubated with DiDesAb-O. In this case, the fibrils had a jagged appearance with the presence of apparently globular structures on their surface (**Figure 3A**). The fibrils appeared again short and thinner than those formed with DesAbO, with a diameter of 7.4 ± 0.6 nm, suggesting that fibrils may be breaking or degrading due to increased structural weakness. Altogether, these findings suggest a more potent inhibitory effect of DiDesAb-O on fibril assembly and maturation (**Figure 3B**).

**Figure 3.**
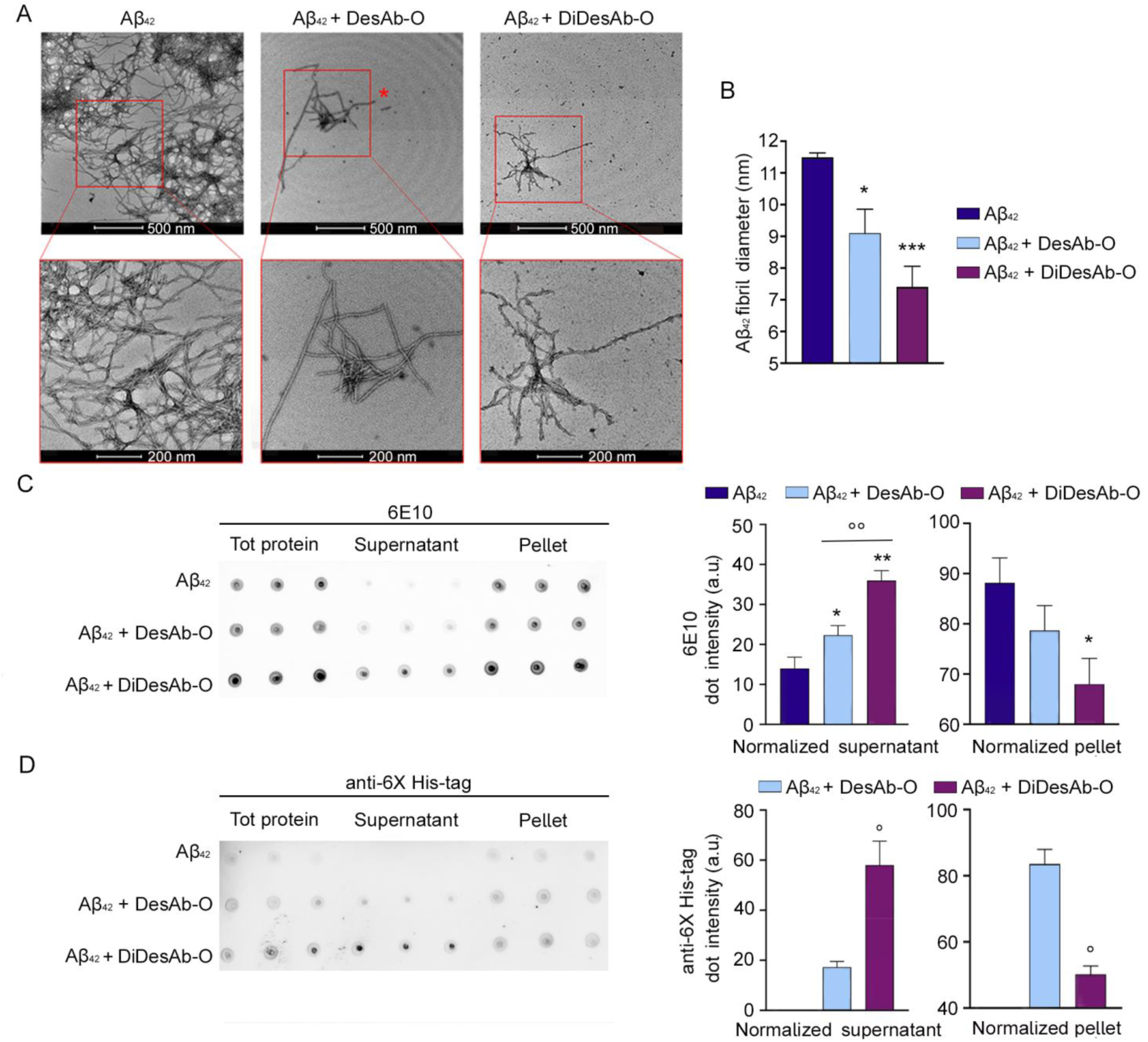
Evaluation of Aβ_42_ fibril fragility and morphological changes induced by co-incubation with sdAbs. **A** TEM images of Aβ_42_ fibrils obtained in the absence or presence of sdAbs after 4 days at 37 °C. Higher magnifications of Aβ_42_ fibrils are showed in the box areas. **B** Aβ_42_ fibrils diameter quantification performed with ImageJ. Error bars are S.E.M (*n* = 3). For each TEM image, 15 to 25 diameter values were obtained and averaged. Diameter values were compared by One-tailed *Student* t-test relative to Aβ_42_ fibrils obtained in absence of sdAbs (*P <0.05 and ***P < 0.001, relative to Aβ_42_ fibrils obtained in the absence of sdAbs). **C,D** Dot Blot of total protein, supernatant and pellet of centrifuged Aβ_42_ fibrils samples obtained with or without co-incubation with sdAbs. The membranes were incubated with 6E10 (**C**) and anti-6X His-tag (**D**) as primary Abs for Aβ_42_ and sdAbs, respectively. Aβ_42_ signal was subtracted as a background. Error bars indicate S.E.M. Statistical analysis was performed by One-tailed *Student* t-test. Samples (*n* = 3 for 6E10 Ab and *n* = 2 for anti-6X His-tag Ab, for both supernatant and pellet) were analysed by comparing the absorbance of Aβ_42_ fibrils obtained without Ab VS Aβ_42_ fibrils obtained in the presence of sdAbs (* *P* < 0.05 VS DesAb-O and ** *P* < 0.01 VS DiDesAb-O, for supernatant and * P < 0.05 VS DiDesAb-O for pellet, respectively) and versus Aβ_42_ fibrils obtained in the presence of DesAb-O compared to Aβ_42_ aggregates obtained in the presence of DiDesAb-O (°° P < 0.01). In panel D, the presence of DesAb-O in the supernatant and in pellet was compared to the presence of DiDesAb-O (° *P* < 0.05 for supernatant and pellet, respectively).

To further investigate the structural integrity of Aβ_42_ fibrils formed with or without sdAbs, we performed a dot blot assay. Following the end of ThT fluorescence kinetic experiments, the plate was incubated at 37°C for 4 days. Samples were then collected and centrifuged to separate the soluble and insoluble protein fractions. Prior to centrifugation, an aliquot of each sample was stored and considered as the total protein amount. To analyze the proportion of soluble and insoluble Aβ_42_ species and Ab fragments, samples were spotted onto a nitrocellulose membrane and incubated with either 6E10 Ab for Aβ_42_ detection or anti-6X His-tag Ab for DesAb-O and DiDesAb-O detection. The signal intensity of the supernatant was normalized to that of the total protein of the corresponding sample. Our results show that Aβ_42_ aggregation in the absence of sdAbs leads to nearly complete conversion of Aβ_42_ monomers into insoluble aggregates, *e.g.*, fibrils, with only 14 ± 3% of protein in the supernatant and 88 ± 5% in the pellet (**Figure 3C**). Conversely, samples obtained in the presence of DesAb-O and DiDesAb-O contained an increased amount of Aβ_42_ in the supernatant (22 ± 2% and 36 ± 2%, respectively) and only partial deposition of Aβ_42_ in the pellet (79 ± 5% and 68 ± 5%, respectively), as shown in **Figure 3C**. These data indicate that the sdAbs, particularly DiDesAb-O, increased the presence of soluble Aβ_42_ species in the aggregation mixture. This result is in agreement with the presence of small globular aggregates in the background of the TEM images, as noted earlier.

Quantification of the sdAb signals showed that DesAb-O was predominantly in the pellet, likely due to its binding to the aggregates, *e.g.*, the oligomers (17 ± 2% and 84 ± 4% for supernatant and pellet, respectively), whereas DiDesAb-O was largely in the supernatant of the samples (58 ± 9% and 50 ± 3% for supernatant and pellet, respectively) (**Figure 3D**). This observation, along with TEM images (**Figure 3A**) and ThT fluorescence time-course analyses (**Figure 2A**), suggests that DiDesAb-O displays stronger binding to and stabilises soluble Aβ_42_ aggregates, *e.g.*, the oligomers, which populate the supernatant to the detriment of fibrils. Alternatively, it is also possible that DiDesAb-O can lead to unstable Aβ_42_ fibrils that more easily can fragment and become soluble.

Finally, we carried out a proteinase K (PK) digestion assay to determine whether the Ab fragments can induce structural modifications in Aβ_42_ fibrils (**Figure S5**). Fibrils were collected from the co-incubation samples with sdAbs at the end of aggregation after 4 days, treated with increasing concentrations of PK followed by Western blotting analysis was performed using the anti-Aβ_42_ N-teminus 6E10 Ab. Aβ_42_ fibrils formed without sdAbs or with DesAb-O appeared more resistant and did not enter the gel, unlike fibrils formed with DiDesAb-O (**Figure S5A**). As far the fractions that enter the gel, band intensities were normalized to those obtained with 0 *μ*g/mL PK in the corresponding samples, taken as 100%. Aβ_42_ fibrils obtained in the presence of DesAb-O displayed an increased resistance to PK digestion at low PK concentrations (10 *μ*g/mL) probably due to conformational changes in the fibrils, making them slightly more resistant to PK catalysed cleavage. In contrast, fibrils obtained in the absence of sdAbs and in the presence of DiDesAb-O followed similar PK resistance (**Figure S5C)**.

Overall, these results demonstrate that DesAb-O and DiDesAb-O induce significant changes in the structure of Aβ_42_ fibrils. As evidenced by dot blot, TEM and Western blotting, DiDesAb-O action results in a reduction of fibril diameter and increased structural fragility.

### DiDesAb-O detects Aβ_42_ oligomers interacting with cellular membranes and internalized into the cytoplasm

Next, we aimed to evaluate the ability of DiDesAb-O to detect Aβ_42_ oligomers in cell cultures and inhibit their toxicity using cell biology experiments. For these experiments, we used synthetic Aβ_42_ rather than the recombinant peptide, as it is endotoxin-free, which is a crucial requirement for these sensitive experiments. First, we ascertained the ability of synthetic Aβ_42_ to form toxic aggregates in the absence of sdAbs in a typical time course experiment (10 µM Aβ_42_ in PBS, pH 7.4, 37 °C) (**Figures S6 and S7**). For the experiments described below with sdAbs, we used the heterogeneous Aβ_42_ aggregate solution formed after 8 h of incubation under the same conditions described above, which we will refer to as “oligomers” as it included mostly oligomeric species that are toxic to the cells. Human neuroblastoma SH-SY5Y cells were exposed to 0.5 μM Aβ_42_ oligomers for 1 h and then treated with sdAbs as primary Abs to detect Aβ_42_ oligomers. Plasma membrane (red channel) and Aβ_42_ oligomers (green channel) were counterstained and analysed by confocal microscopy (**Figure 4A**), as previously reported (Bigi, Napolitano et al., 2024). At 3 μM, both DiDesAb-O and DesAb-O caused a green fluorescence increase in cells treated with oligomers by 205 ± 18% and 172 ± 13%, respectively, as compared to cells treated with sdAbs but not oligomers (**Figure 4B**). At 1 μM, only DiDesAb-O was able to produce a significant increase of green fluorescence (173 ± 14% for DiDesAb-O vs 140 ± 17% for DesAb-O), indicating greater binding to oligomers (**Figure 4B**). 6E10 Ab, that was used as a control, can recognize both low and high molecular weight structures, as revealed by the presence of green dots with different size (**Figure 4A**). On the contrary, both sdAbs can detect only small oligomeric species within the heterogeneous mixture, representing further evidence of their specificity for Aβ_42_ oligomeric species. Overall, in our experimental conditions, DiDesAb-O selectively recognized toxic Aβ_42_ oligomers interacting with cellular membranes and penetrated into the cytosol even at lower concentrations than DesAb-O, suggesting a stronger binding.

**Figure 4.**
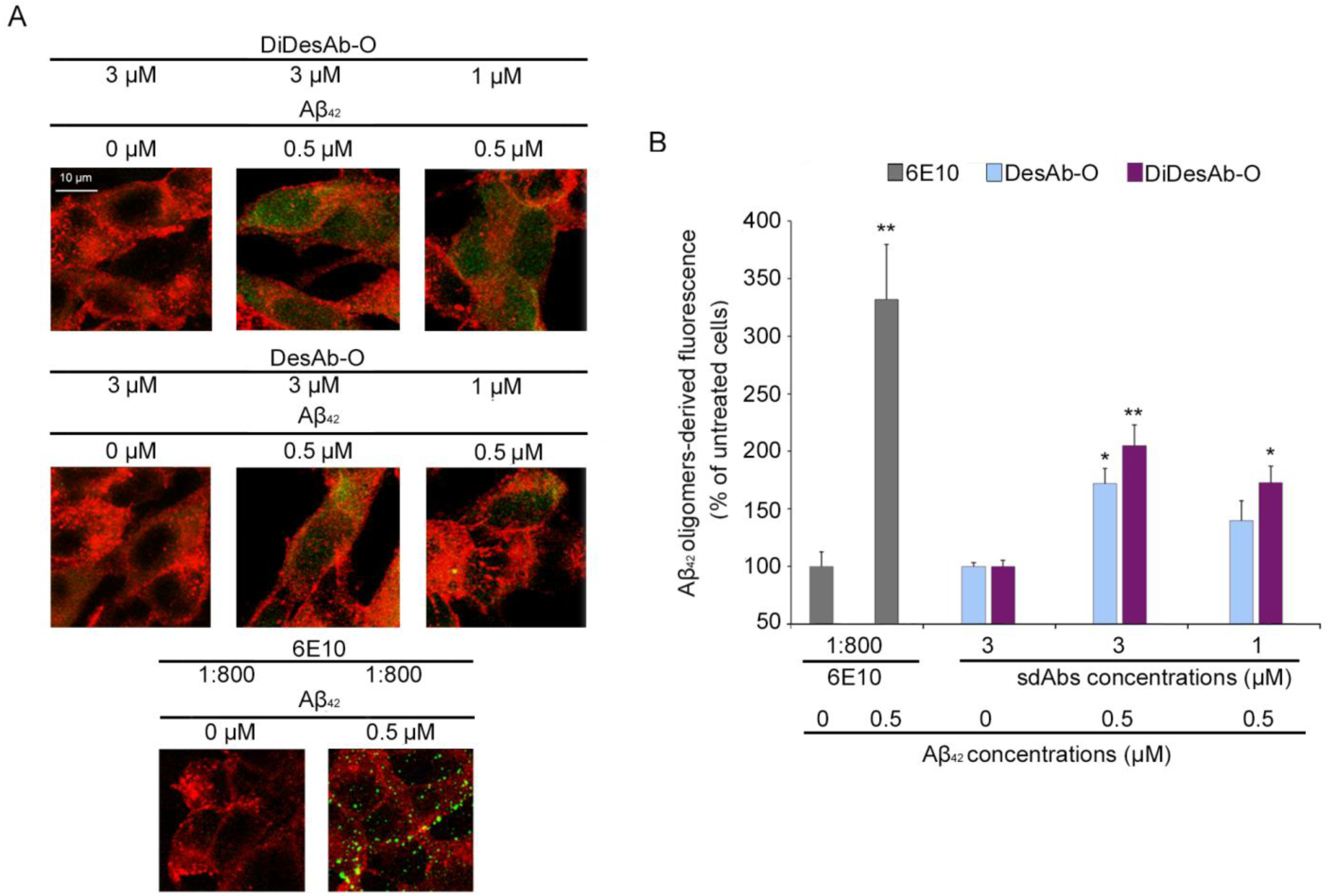
DiDesAb-O detects Aβ_42_ oligomers interacting with neuronal cells and penetrated into the cytosol at lower concentrations than DesAb-O. **A** Representative confocal microscopy images of SH-SY5Y cells left untreated or treated with Aβ_42_ species at 0.5 μM for 1 h. Red and green fluorescence indicates the cell membranes and the Aβ_42_ oligomers, respectively, detected with wheat germ agglutinin (WGA) and sdAbs at two different concentrations (3 μM or 1 μM), respectively. **B** The bar plot represents the results of a semi-quantitative analysis of green fluorescence. Error bars indicate S.E.M. Statistical analysis was performed by ANOVA with multiple comparison. Sample (*n* = 3) were analysed by One-way ANOVA followed by Bonferroni’s multiple-comparison test to their respective untreated cells (\**P* < 0.05 and \*\**P* < 0.01). 200–250 cells were analyzed per condition.

### DiDesAb-O inhibits the interaction of Aβ_42_ oligomers with neuronal membranes

We evaluated whether DiDesAb-O was able to neutralize Aβ_42_ oligomers, preventing their interaction with neuronal membranes. To do so, 0.5 μM Aβ_42_ oligomers formed after 8 h of incubation were incubated for 1 h with DesAb-O and DiDesAb-O at varying molar ratios Aβ_42_:sdAbs (1:0.1, 1:0.25, 1:0.5, 1:1, 1:2, 1:3). These solutions were then added to the cell culture medium of SH-SY5Y cells for 15 min, as previously reported (Bigi, Napolitano et al., 2024). To detect only the oligomers bound to the cell surface, the cellular membrane was not permeabilized at this stage, thus preventing oligomer internalization. The binding of the aggregates on the cellular membranes was assessed by confocal microscopy using the 6E10 Ab as a probe. Aβ_42_ oligomers colocalized with cellular membranes in the absence of pre-incubation with the sdAbs (**Figure 5A**), as previously reported (Bigi et al., 2020; Bigi, Napolitano et al., 2024). Following the pre-incubation step, the interaction between Aβ_42_ oligomers and the neuronal membranes was significantly reduced (**Figure 5A**). In particular, DiDesAb-O appeared to be more efficient than DesAb-O, being more effective at all sdAb concentrations tested and producing the same inhibition as DesAb-O at 10-fold lower concentrations (**Figure 5B**). These results suggest DiDesAb-O binds stronger to Aβ_42_ oligomers with respect to DesAb-O.

**Figure 5.**
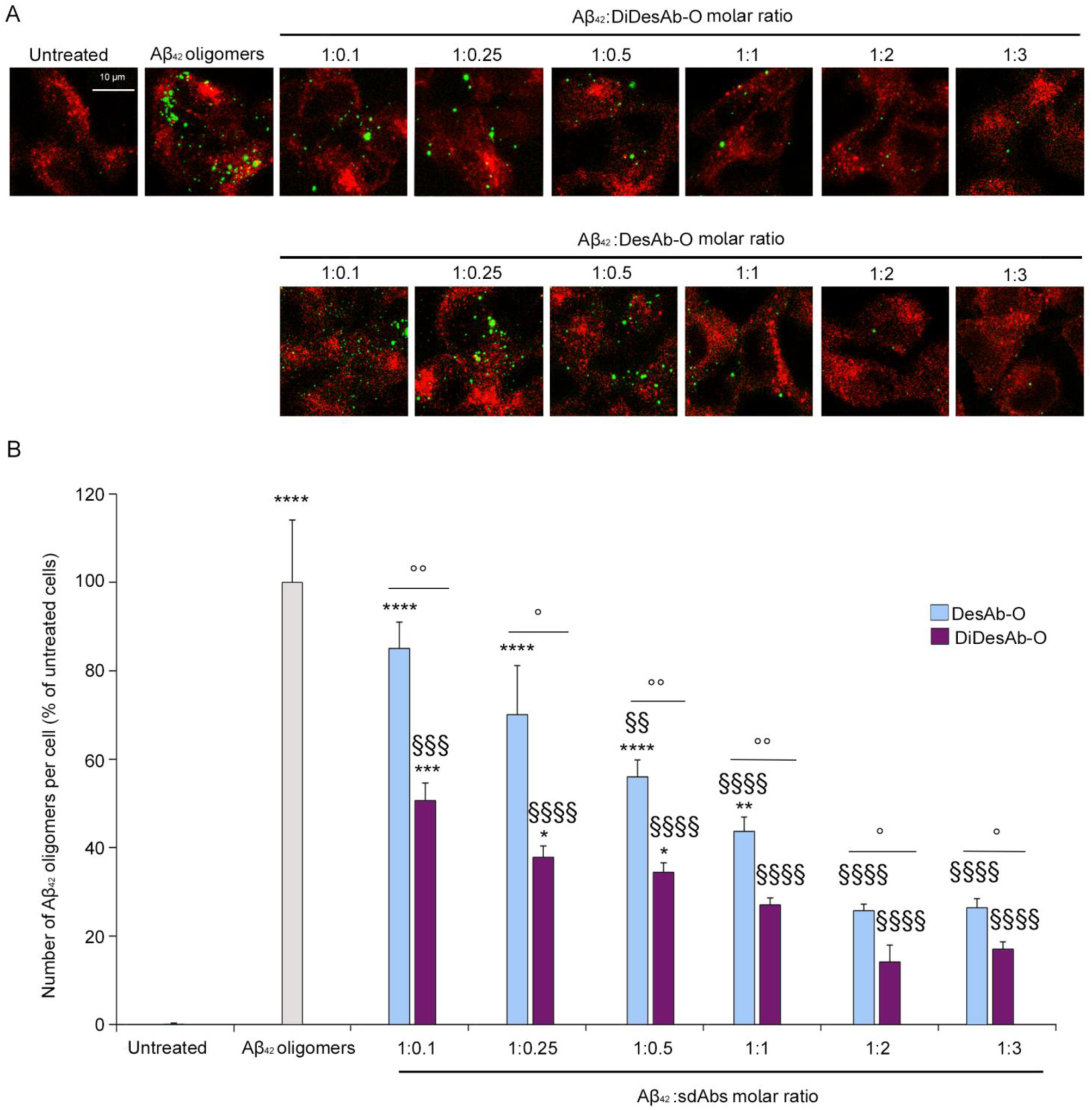
DiDesAb-O inhibits the binding of Aβ_42_ oligomers to the neuronal membrane to a greater extent with respect to DesAb-O. **A** Representative confocal microscopy images of SH-SY5Y cells treated with 0.5 μM Aβ_42_ oligomers following 1 h pre-incubation in the absence (first row) or presence of DiDesAb-O (second row) or DesAb-O (third row) at the indicated Aβ_42_:Abs molar ratios. Red and green fluorescence indicates the cell membranes and Aβ_42_ oligomers detected with WGA and 6E10 Ab, respectively. **C** Counting of Aβ_42_ oligomers bound to the cellular membrane measured following incubation under the conditions represented in panels A. Error bars indicate S.E.M. Statistical analysis was performed by ANOVA with multiple comparison or by *Student* t-test. Samples (*n* = 3) were analysed by One-way ANOVA followed by Bonferroni’s multiple-comparison test to untreated cells (** *P* < 0.05, ** *P* < 0.01, *** *P* < 0.001 and **** *P* < 0.0001), or to cells treated with Aβ_42_ oligomers only (§§ P < 0.01 and §§§§ *P* < 0.0001). sdAbs at the molar ratio were analysed by One-tailed *Student* t-test (° P < 0.05 and °° P < 0.01).

### DiDesAb-O prevents Aβ_42_ oligomer toxicity

We then evaluated the ability of DiDesAb-O to prevent critical detrimental effects induced by Aβ_42_ oligomers. First, we monitored the disruption of cytosolic Ca^2+^ homeostasis induced by Aβ_42_ oligomers, as previously reported (Cascella et al., 2021; Bigi, Napolitano et al., 2024). SH-SY5Y cells were treated for 15 min with solutions containing Aβ_42_ oligomers and sdAbs at varying molar ratios. Aβ_42_ oligomers without sdAbs caused an extensive Ca^2+^ influx, with an increase of 280 ± 2% relative to untreated cells (**Figure 6A,B**). Pre-incubation of oligomers with DiDesAb-O was found to generate a significant protective effect at 6-fold lower concentration than DesAb-O (**Figure 6A,B**). DiDesAb-O was protective even at the lowest concentration tested (0.05 µM, 1:0.1 ratio). Of note, neither sdAb affects neuronal viability when added alone to the cell medium (**Figure 6A,B**).

**Figure 6.**
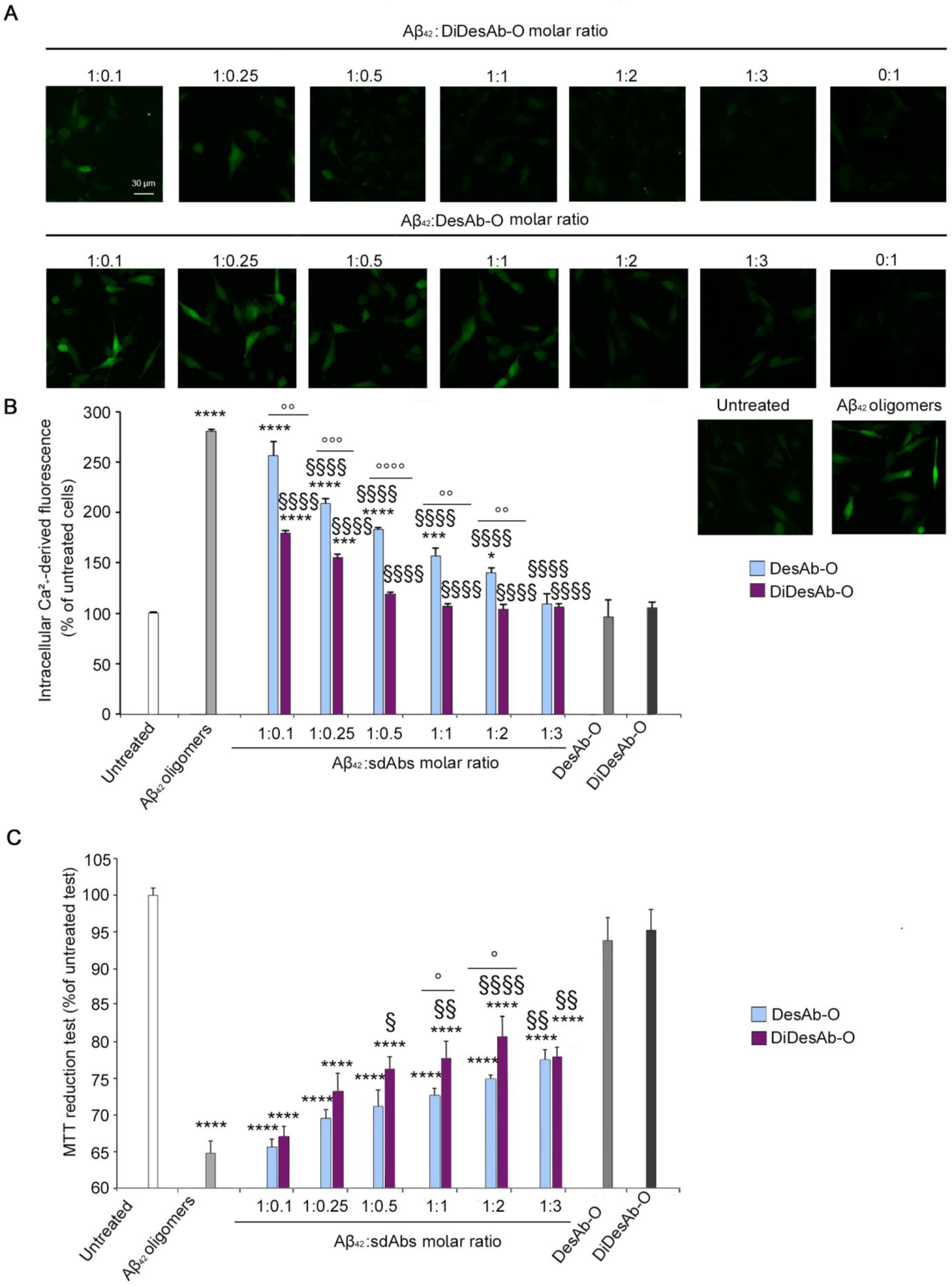
DiDesAb-O strongly prevents the neurotoxicity induced by Aβ_42_ oligomers. **A** Representative confocal microscopy images showing the Ca^2+^-derived fluorescence in SH-SY5Y cells treated for 15 min with 0.5 µM Aβ_42_ oligomers pre-incubated with or without increasing molar ratios (1:0.1, 1:0.25, 1:0.5, 1:1, 1:2, 1:3) of DiDesAb-O (**top**) or DesAb-O (**bottom**). Cells were then loaded with the Fluo-4AM probe as described in the Materials and Methods section. **B** Semi-quantitative analyses of the Ca^2+^-derived fluorescence expressed as the percentage of the value for untreated cells, taken as 100%. **C** MTT reduction in SH-SY5Y cells treated for 24 h with increasing Aβ_42_ oligomers:sdAbs molar ratios (1:0.1, 1:0.25, 1:0.5, 1:1, 1:2, 1:3). Values are expressed as the percentage of the value for untreated cells, taken as 100%. **B, C** Error bars indicate S.E.M. Statistical analysis was performed by ANOVA with multiple comparison or by *Student*-t test. Samples (*n* = 3 for confocal experiment and *n* = 4 for MTT) were analysed by One-way ANOVA followed by Bonferroni’s multiple-comparison test to untreated cells (* *P* < 0.05, *** *P <* 0.001 and **** *P* < 0.0001), or to cells treated with Aβ_42_ oligomers (§ P < 0.05, §§ P < 0.01 and §§§§ *P* < 0.0001). sdAbs at the same molar ratio were analysed by One-tailed *Student* t-test (° *P* < 0.05, °° *P* < 0.01, °°° *P* < 0.001 and °°°° *P* < 0.0001).

The protective effect of DiDesAb-O was also evaluated by analyzing, with the MTT reduction test, the mitochondrial status of cultured cells treated with Aβ_42_ oligomers (**Figure 6C**). Aβ_42_ oligomeric species at 0.5 μM were incubated in the absence or presence of varying concentrations of sdAbs for 1 h and then added to the culture medium of SH-SY5Y cells for 24 h. While Aβ_42_ oligomers significantly reduced (by 35 ± 2%) the mitochondrial activity of SH-SY5Y cells as compared to untreated cells, as previously reported (Bigi, Napolitano et al., 2024), pre- incubated with the sdAbs allowed a significant recovery of the mitochondrial functionality (**Figure 6C**). Again, DiDesAb-O was more effective than DesAb-O at most Ab concentrations tested. sdAbs alone were, by contrast, ineffective (**Figure 6C**).

Considering these results together, we can confirm the increased ability of DiDesAb-O to prevent Aβ_42_ oligomer-induced toxicity.

### DiDesAb-O prevents toxic effects induced by the CSFs of AD patients

Following the previously reported ability of DesAb-O to selectively detect Aβ_42_ oligomers in cultured cells exposed to CSFs of AD patients (Bigi, Napolitano et al., 2024), we performed a proof-of-concept experiment on a small set of clinical samples of CSF (n = 4 from AD patients and n = 4 from controls subjects) to evaluate whether DiDesAb-O was able to neutralize the cytotoxicity induced by Aβ_42_ oligomeric species present in AD CSFs and whether the engineering of the DiDesAb-O sdAb into a dimeric form resulted in improved performance (**Figure 7A,B**). Thus, we monitored the dysregulation of cytosolic Ca^2+^ homeostasis in SH-SY5Y treated for 5 h with the CSFs from AD patients and control subjects diluted 1:1 with the cell culture medium, in the absence or presence of a 1 h pre-incubation with 3 μM or 1 μM sdAbs, as previously reported for the monomeric sdAb (Bigi, Napolitano et al., 2024). The AD CSFs significantly increased the intracellular Ca^2+^ concentration by 190 ± 7% with respect to untreated cells, whereas control CSFs increased it by only 137 ± 6% (**Figure 7A,B**). After 1 h of pre-incubation with 3 μM, both DesAb-O and DiDesAb-O completely reduced the Ca^2+^ dyshomeostasis induced by AD CSFs (93 ± 3% and 84 ± 5% relative to untreated cells, respectively) and control CSFs (96 ± 5% and 95 ± 6%, relative to untreated cells, respectively) (**Figure 7B**). After 1 h of pre-incubation with 1 μM, DesAb-O lost in part its efficacy, whereas DiDesAb-O continued to be active against AD CSFs (**Figure 7B**).

**Figure 7.**
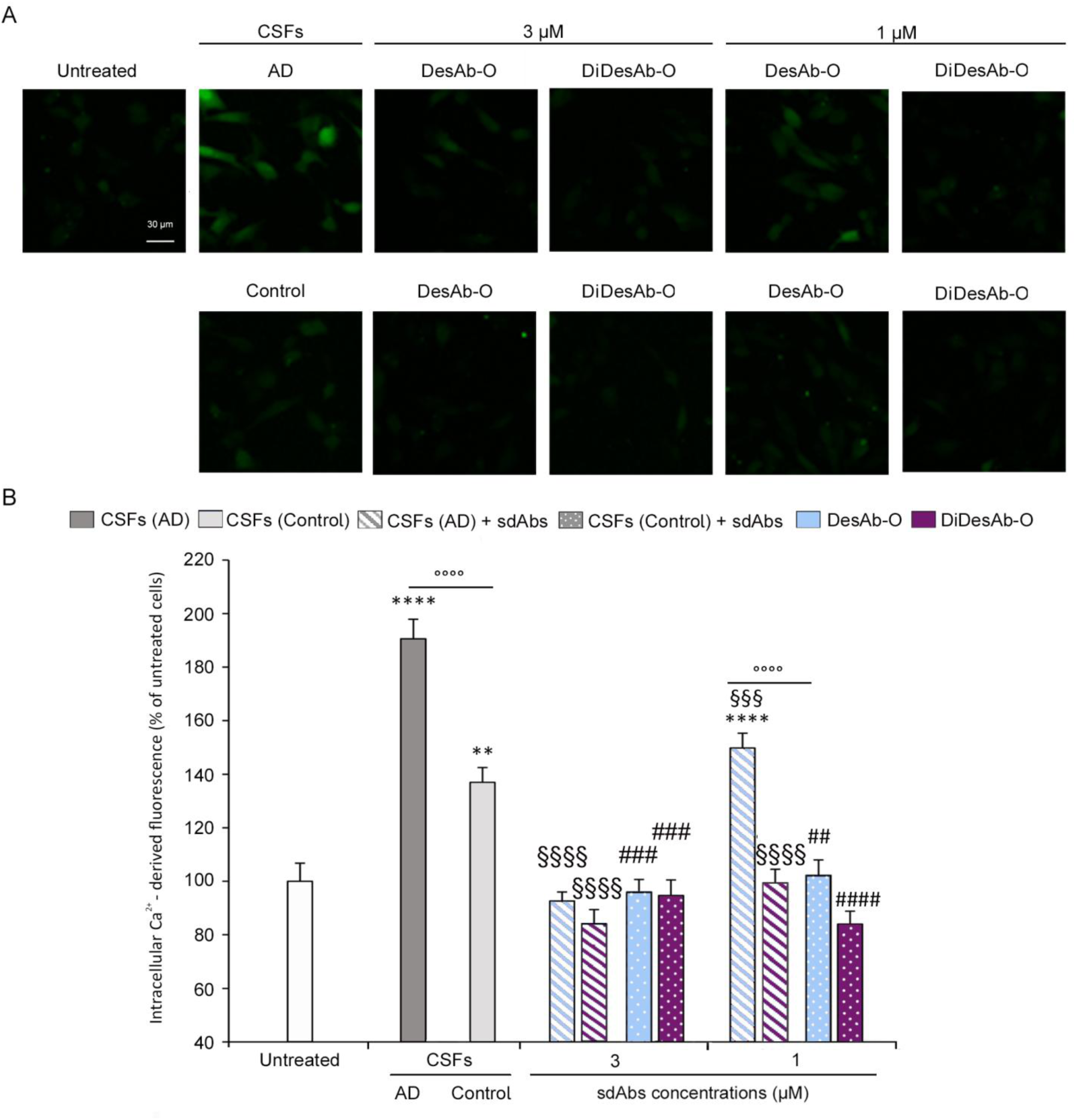
DiDesAb-O significantly reduce the Ca^2+^ dyshomeostasis induced by CSFs derived from AD patients. Intracellular Ca^2+-^derived fluorescence in SH-SY5Y cells treated for 5 h with CSFs from AD patients (stripped bars) and age-matched control subjects (dotted bars) (n = 4 for AD and n = 4 for control subjects), diluted 1:1 with the extracellular medium, following 1 h pre-incubation in the absence (grey bars) or presence of DesAb-O (light blue bars) and DiDesAb-O (purple bars) at 3 μM or 1 μM, as indicated. Error bars indicate S.E.M. Statistical analysis was performed by ANOVA with multiple comparison. Samples were analysed by One-way ANOVA followed by Bonferroni’s multiple-comparison test relative to untreated cells (**P < 0.01 and ****P < 0.0001) or to cells treated with AD CSF samples without sdAbs (§§§ P < 0.001 and §§§§ P < 0.0001), or to cells treated with control CSF samples without sdAbs (## P < 0.01, ### P < 0.001 and #### P < 0.0001) or to compare AD and controls CSF samples under the same conditions (°°°° P < 0.0001).

Taken together, these data demonstrate the ability of DiDesAb-O to selectively neutralize toxic species present in the CSFs of AD patients, to an extent larger than that of the monomeric counterpart.

## Discussion

Over the past two decades, several mAbs have been developed to detect or remove Aβ aggregates in the brains of AD patients (Karran and De Strooper, 2022). Some of these, such as Lecanemab and Donanemab, have received FDA approval (van Dyck et al., 2023; Sims et al., 2023). Despite their clinical potential, many trials have failed and even the FDA-approved mAbs are controversial due to their severe side effects (van Dyck et al., 2023; Sims et al. 2023; Cummings et al., 2024). The large molecular size of conventional mAbs (∼150 kDa) significantly limits their ability to cross the BBB, as only a small fraction (typically less than 0.1%) reaches the CNS. This low bioavailability reduces their efficacy in targeting Aβ aggregates in the brain. Furthermore, mAbs have a significant immunogenicity, and off-target binding, potentially leading to side effects.

Ab fragments, such as sdAbs, represent highly promising biomolecules for diagnostic and therapeutic applications for a range of neurological disorders, including dementia. They offer several advantages over mAbs including high solubility, low immunogenicity (Lafaye et al., 2009; Muyldermans, 2013; Ackaert et al., 2021; De Leiris et al., 2024). Furthermore, sdAbs can be engineered with a variety of protein engineering approaches to enhance different properties, such as the BBB penetration (Wouters et al., 2020; Ruiz-López and Schuhmacher, 2021; Yogi et al., 2022) and microglia activation for degradation of neurotoxic aggregates from brain parenchyma (Zhao et al., 2022). To date, numerous Ab fragments have been developed against various types of Aβ aggregates, showing promising results in preclinical studies by neutralizing Aβ aggregate toxicity (Zameer et al., 2008; Lafaye et al., 2009; Kasturirangan et al., 2012; Nisbet et al., 2013; Zheng et al., 2022). Beyond therapeutic applications, sdAbs such as R3VQ and A2 are effective *in vivo* imaging agents due to their ability to bind Aβ and tau aggregates (Li et al., 2016). While all these molecules are promising, they often recognize non-toxic monomers and/or less-toxic amyloid fibrils (Aprile et al., 2020). Thus, higher specificity for the oligomers is required to enable clinical applications. Additionally, most of these molecules are derived from non-human framework sequences, which may trigger the formation of anti-drug Abs (ADA) when administered to humans regularly (Rossotti et al., 2022). Thus, Ab fragments consisting of human-derived Ab domains may find more streamlined clinical applications.

Building on the promising functional properties of the previously characterized sdAb, DesAb-O (Aprile et al., 2020; Bigi, Napolitano et al., 2024), we sought to enhance its avidity and binding ability for Aβ_42_ oligomers by designing an improved dimeric version, named DiDesAb-O. Our hypothesis was that Aβ_42_ oligomers present multiple copies of the same target epitope. By making DesAb-O dimeric, it could simultaneously bind to multiple epitopes, thereby increasing binding strength through avidity. The use of flexible linkers to engineer sdAbs for enhanced binding avidity and specificity has been successfully demonstrated (Hultberg et al., 2011; Cardoso et al., 2014). In our approach, we created a dimeric structure by linking two monomers with a (GGGGS)₃ linker.

Our functional analyses revealed that DiDesAb-O displays stronger oligomer binding and anti-aggregation activity compared to monomeric DesAb-O. Furthermore, it can induce drastic morphological changes in Aβ_42_ fibrils. Other Ab fragments specific for Aβ_42_ oligomers, such as V31-1 (Lafaye et al., 2009), A4-scFv, and E1 (Kasturirangan et al., 2012), have been shown to inhibit fibril formation. Specifically, Dynamic Light Scattering (DLS) showed an inhibitory effect of V31-1 (Lafaye et al., 2009) whereases AFM analysis confirmed the same property for A4-scFv and E1 (Kasturirangan et al., 2012).

We also showed the ability of DiDesAb-O to selectively detect Aβ_42_ oligomers in cultured cells, following the approach of our previous work with DesAb-O (Bigi, Napolitano et al., 2024). We found that pre-incubation of Aβ_42_ oligomers with DiDesAb-O resulted in a significant reduction of their interaction with neuronal membranes, more effectively than with DesAb-O. Furthermore, we observed reduced oligomer-induced intracellular Ca^2+^ flux and improved prevention of the mitochondrial dysfunction compared to DesAb-O, suggesting an enhanced detection of key epitopes in the structure of the toxic oligomers.

Prior to this work, other mAbs (Lambert et al., 2007; He et al., 2021) or Ab fragments (Nisbet et al., 2013; Bitencourt et al., 2020; Haynes et al., 2024) demonstrated their ability to counteract Aβ_42_ oligomer-induced toxicity. However, from all these studies it is difficult to extrapolate the exact type of aggregate species in the milieu of metastable species recognized by Abs or Ab fragment. At present, only V31-1 (Lafaye et al., 2009), A4-scFv, E1 (Kasturirangan et al., 2012), DesAb-O (Aprile et al., 2020) and DiDesAb-O studied here, target specifically Aβ_42_ oligomers. Given that targeting specific conformational Aβ species can lead to markedly different outcomes, our study focuses on enhancing the binding ability of DesAb-O to more effectively target toxic oligomers. Improving oligomer specificity may not only increase therapeutic efficacy but also allow for lower dosage requirements, potentially minimizing the risk of adverse effects.

To evaluate the improved specificity in a biologically relevant context, we conducted further analyses to determine whether DiDesAb-O could specifically bind to neurotoxic Aβ_42_ oligomers present in AD CSFs and effectively neutralize their toxic effects on neuroblastoma cells. DiDesAb-O was shown to prevent neuronal dysfunction caused by Aβ_42_ oligomers in the CSFs of AD patients at lower concentrations than DesAb-O. This finding highlights its enhanced sensitivity for Aβ_42_ oligomers and its capability to detect these toxic species within complex biological fluids, such as AD CSF.

## Conclusions

This study demonstrates that engineering sdAbs and, in particular, rational engineering of dimeric sdAbs, can significantly enhance their binding to specific protein species, *e.g.*, Aβ_42_ oligomers. We developed DiDesAb-O, a dimeric version of the single-domain antibody DesAb-O, which exhibits improved binding to toxic amyloid-β oligomers both *in vitro* and in the CSF of AD patients and greater efficacy in inhibiting their toxicity. While further validation in larger patient cohorts is needed, these findings provide a robust foundation for creating next-generation Ab fragments with enhanced binding to heterogeneous protein aggregates, opening new avenues for earlier diagnosis, earlier intervention and better disease management for AD and other neurodegenerative diseases.

## Supporting information

DiDesAb-O Supplementary text

## Declarations

### Ethics approval and consent to participate

Not applicable

### Consent for publication

Not applicable

### Availability of data and materials

The authors confirm that all data needed to evaluate the conclusions of this study are available within the article.

### Author Contributions

LN: investigation, validation, formal analysis, data curation, visualisation, writing—original draft, writing—review and editing. DMV: investigation, validation, formal analysis, data curation, writing—original draft, writing—review and editing. AB: investigation, validation, formal analysis, data curation, visualisation, writing—original draft, writing—review and editing. CC:, funding acquisition, writing—review and editing, supervision. FC: conceptualization, methodology, formal analysis, writing—original draft, writing—review and editing, supervision, funding acquisition. RC: conceptualization, methodology, formal analysis, writing—original draft, writing—review and editing, supervision, project administration, funding acquisition. FAA: conceptualization, methodology, formal analysis, writing—original draft, writing—review and editing, supervision, project administration, funding acquisition.

