## Supplementary material for "Engineering a Dimeric Single-Domain Antibody for Improved Detection and neutralization of Amyloid-β Oligomers": DiDesAb-O Supplementary text

### Supplementary materials and methods

**Protein Expression and Purification.** The DiDesAb-O construct was expressed and purified using a pET28a (+) vector in *E. coli* Origami 2 (DE3) Competent Cells (Merck Millipore), as previously described (Sormanni et al, 2015). Cells were grown for 24 h at 37 °C using LB medium (Merck Millipore) supplemented with Kanamycin (50 μg /ml). Cells were harvested by centrifugation, resuspended with 20 mM phosphate buffer, pH 8.0, with the addition of one EDTA-Free Complete Protease Inhibitor Cocktail Tablet (Roche) per 500 ml of cell growth, and lysed using sonication. Cell debris was removed using centrifugation at 48,400 g (JA-20 rotor, Beckman Coulter) for 45 min. The cleared supernatant lysate was loaded onto a $Ni^{2+}$-NTA HisTrap Superflow column (Cytiva), previously equilibrated with 20 mM phosphate buffer, pH 8.0, containing 10 mM imidazole. After washing with 20 mM phosphate buffer containing 40 mM imidazole, the His-tagged DiDesAb-O was eluted with 20 mM phosphate buffer containing 200 mM imidazole and dialyzed extensively against 20 mM phosphate buffer. DiDesAb-O was finally purified using size-exclusion chromatography with a HiLoad 16/600 Superdex 75 pg column (GE Healthcare), previously equilibrated in 20 mM phosphate buffer, pH 8.0. The protein concentration was determined by absorbance measurement at 280 nm using theoretical extinction coefficients calculated with ExPASy ProtParam (Gill and Von Hippel, 1989). Both the flow through and peak fractions were then loaded on 4−12% Bis-Tris NuPAGE gels (Thermo Fisher Scientific) to verify the sample purity.

Exression and purification of $A\beta_{42}$ was carried out as previously described (Abelein et al, 2020; Vadukul et al., 2023). Aliquots of purified $A\beta_{42}$ were stored at −80 °C. For all cellular biology experiments we purchased synthetic $A\beta_{42}$ peptides from Bachem to obtain amyloid β-derived diffusible ligands (ADDLs) oligomers (Lambert et al., 1998).

**Circular Dichroism**. Far-ultraviolet (far-UV) CD spectra of DiDesAb-O were acquired using a Chirascan spectropolarimeter (Applied Photophysics) equipped with a Peltier temperature control unit. Measurements were performed in a quartz cuvette with 1 mm path length. Samples contained 6 μM protein in 10 mM $Na_2HPO_4$, 1.8 mM $KH_2PO_4$, 137 mM NaCl and 2.7 mM KCl, pH 7.4 (phosphate-buffered saline or PBS). The far-UV CD spectra of DiDesAb-O were recorded from 200 to 240 nm at 20 °C, and the spectrum of the buffer was subtracted from the spectra of DiDesAb-O. Accumulation was 10. In another set of experiments, CD spectra of 6 μM DesAb-O or DiDesAb-O were recorded between 20 °C and 90 °C with 5 °C intervals. A background spectrum of the sample buffer was subtracted from all sample spectra. Raw data of θ (units of mdeg) were converted to mean residue ellipticity ($[\theta]_{res}$, units deg $cm^2$ $dmol^{-1}$) using (Greenfield, 2006):

$$[\theta]_{res} = \theta / [(n-1) \bullet l \bullet c] \quad (2)$$

where $[\theta]_{res}$, is in deg $cm^2$ $dmol^{-1}$, θ is in mdeg, n is the number of amino acid residues, l is the cuvette pathlength in mm, and c is the protein concentration in M. The denaturation curves were obtained by plotting $[\theta]_{res}$ against temperature, fitted with the Santoro and Bolen equation and normalised to fraction folded (%) values:

$$[\theta]_{res} = \frac{[([\theta]_{res}(F) + m(F) * T) + ([\theta]_{res}(U) + m(U) * T) * e^{\left(\frac{-\Delta G}{RT}\right)}]}{[1 + e^{\left(\frac{-\Delta G}{RT}\right)}]} \quad (3)$$

where $[\theta]_{res}$ is the measured molar residue ellipticity at temperature $T$ (°C), $[\theta]_{res}(F)$ and $[\theta]_{res}(U)$ are the $[\theta]_{res}$ values for the folded and unfolded states at 20 °C, respectively, $m(F)$ and $m(U)$ are the slopes of the folded and unfolded baselines, respectively, $\Delta G$ is the Gibbs free energy change upon unfolding and $R$ is the universal gas constant.

By fitting the data obtained with the above equation, it was possible to determine the $\Delta G$ with the temperature increase. In order to calculate the temperature of half-denaturation ($T_m$), we firstly determined the fraction folded with the following equation:

$$Fraction\ folded\ (\%) = \frac{([\theta]_{res} - [\theta]_{res}(U))}{([\theta]_{res}(F) - [\theta]_{res}(U))}$$

The $T_m$ values represented the temperature at which the protein is 50% folded and 50% unfolded (fraction folded = 50%).

**Electrospray Ionization Mass Spectrometry.** Purified DiDesAb-O (~20 µM) was analysed by electrospray ionization mass spectrometry (ESI-MS) to confirm molecular weight and sample purity. ESI-MS was performed by the Chemistry Mass Spectrometry facility at the Molecular Sciences Research Hub, Imperial College London.

**PK Digestion and Western Blotting**. Fibrils of recombinant A$\beta_{42}$ (5 µM) obtained after 4 days at 37 °C under constant conditions in the absence or in the presence of sdAbs were centrifuged at max speed (~17,000g) for 1 h and the supernatant was discarded. The pellet was resuspended in 20 mM phosphate buffer and treated with increasing proteinase K (PK) concentrations (0, 10, 25, 50 µg/ml) for 30 min at RT. Samples were then incubated at 95 °C for 5 min to stop the enzymatic reaction, loaded on 4−12% Bis-Tris NuPAGE gels and transferred onto a 0.45 µm nitrocellulose membrane for 7 min at 20 V with the iBlot 2 (Thermo Fisher Scientific). Blocking, incubation with 1:1000 diluted 6E10 Ab in 0.1% PBS-Tween, and detections were carried out as described previously (Vadukul et al., 2023). Data analysis was performed setting the band intensity at 0 µg/ml PK of each sample as the 100%. Gross values of band intensities of 0 µg/ml PK samples were compared as well.

**Thioflavin T Fluorescence Assays**. To perform *in vitro* aggregation assays for cell biological experiments, the lyophilized synthetic peptide (Bachem) was resuspended in PBS, pH 7.4, resulting in a final concentration of 10 µM. For visualization of the emerging β-sheets in A$\beta_{42}$, samples were added with a final concentration of 25 µM ThT, gently vortexed and pipetted into non binding surface black 96-well plates (Grenier Bio-One) in quadruplets. The plate was read in a BioTek Synergy$^{TM}$ H1 Hybrid Multi-mode reader (Agilent) at 37 °C. The excitation and emission wavelengths were set to 440 and 485 nm, respectively. Buffer-only values were not subtracted from the sample readings but shown in the final graph. Readings were taken every 2 min. The data were plotted using GraphPad Prism version 9.3.1 for Windows (GraphPad Software). To characterize the different type of aggregates formed during the A$\beta_{42}$ aggregation process, we collected A$\beta_{42}$ samples at various timepoints (0, 2, 4, 8, and 24 h) to conduct further experiments (see details below on the subsections of S6 and S7).

**Dot Blot**. To characterize the different types of A$\beta_{42}$ aggregates formed during the aggregation process of the synthetic peptide (Bachem), 2.0 µl (equivalent to 0.1 µg) of each sample were collected at five different timepoints (0, 2, 4, 8, and 24 h) and spotted onto a nitrocellulose membrane. Following a 45-min blocking step (1.0% bovine serum albumin, BSA, in TBS-Tween 0.1%), the membrane was incubated with 1:15.000 diluted human monoclonal anti-ADDLs (19.3) Ab (Creative Biolabs), 1:1000 diluted rabbit polyclonal anti-amyloid fibrils (OC) Ab (Sigma-Aldrich) and 1:800 6E10 Ab for 1 h and 30 min. Subsequently, the membrane was washed three times in TBS-Tween 0.1% for 10 min each and incubated with 1:3000 diluted goat anti-6X His tag (Abcam), goat anti-human (Sigma-Aldrich), or goat anti-rabbit (Abcam) or rabbit anti-mouse

(Abcam) Abs, all conjugated with horseradish peroxidase (HRP) for 1 h. After three additional washes in TBS-Tween 0.1%, the immunolabelled dots were detected using a Super-SignalWest Dura (Pierce) ImageQuant™ TL software (GE Healthcare UK Limited version 8.2) as previously reported (Bigi and Napolitano et al., 2024).

**STED microscopy**. Synthetic A$\beta_{42}$ aggregates collected at different timepoints (0, 2, 4, 8, and 24 h) were spotted on a glass coverslip for 30 min. Then, the samples were blocked with 1x casein for 30 min, washed with PBS, and incubated with 1:500 diluted 19.3, 1:800 diluted 6E10 or 1:800 diluted OC primary Abs for 1 h and 30 min. After washing with PBS, the samples were incubated with 1:500 diluted AF488-conjugated anti-human, AF514-conjugated anti-mouse and AF514-conjugated anti-rabbit secondary Abs. Fluoromount-G™ (Fisher Scientific) was used as mounting medium. STED xyz images (i.e., z-stacks were acquired as previously reported (Bigi A et al., 2024) at 0.1 μm intervals along 3 directions: x, y, and z axes) in bidirectional mode with a Leica SP8 STED 3X confocal microscope system equipped with a Leica HC PL APO CS2 100x/1.40 oil STED White objective. AF514 was excited with a 510 nm-tuned WLL and emission collected from 532 to 551 nm. Frame sequential acquisition was applied to avoid fluorescence overlap. A gating between 0.3 to 6 ns to avoid collection of reflection and autofluorescence was applied. 650 nm pulsed depletion laser was used for AF514 excitation. Collected images were de-convolved with Huygens Professional software version 18.04 (Scientific Volume Imaging B.V.) and analyzed with Leica Application Suite X (LAS X) software (Leica).

**MTT reduction assay.** To assess the cytotoxicity of synthetic A$\beta_{42}$ aggregates formed during the A$\beta_{42}$ aggregation process, a 3-(4,5-dimethyltohiazol-2-yl)-2,5-diphenyltetrazolium bromide (MTT) reduction assay was conducted. 1 μM A$\beta_{42}$ species (monomer equivalents) collected from a ThT aggregation assay at different timepoints (0, 2, 4, 8 and 24 h) were added for 24 h to culture medium of SH-SY5Y cells. 1 μM A$\beta_{42}$ ADDLs were used as positive control. For many experiments, we used A$\beta_{42}$ oligomers obtained after 8 h of incubation at 37 °C. We also performed an MTT test with decreasing concentrations 8 h A$\beta_{42}$ oligomers (1 μM, 0.5 μM, 0.25 μM, 1 nM, 0.5 nM, 0.25 nM and 1 pM) to optimize our experimental conditions. MTT test were performed as reported in the main text.

### Supplementary figures and legends

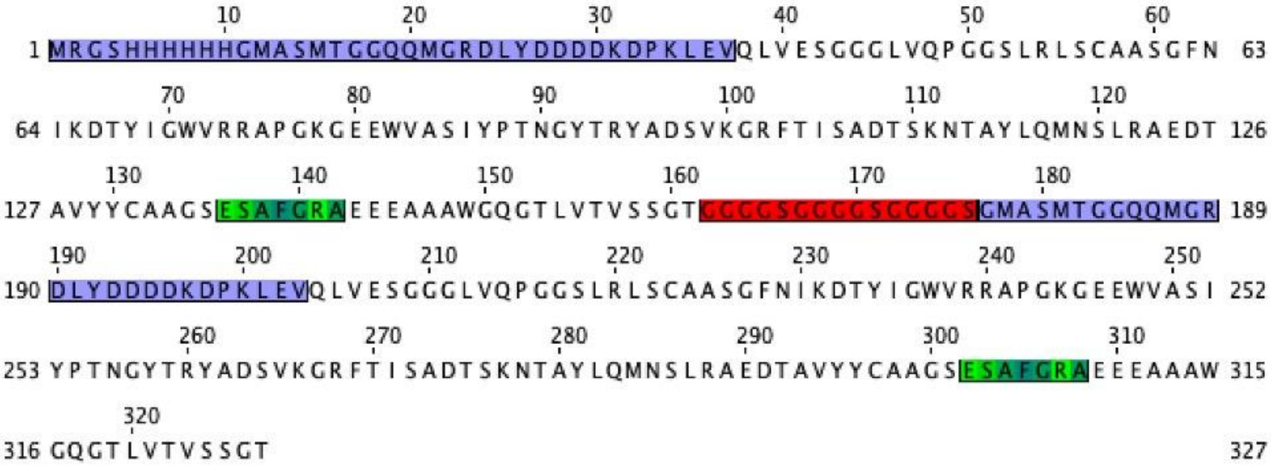


**Figure S1. DiDesAb-O amino acid residues sequence.** N-terminus are highlighted in light purple, the flexible linker $(GGGGS)_3$ is highlighted in red, whereas the antibody binding sites are highlighted in shades of green. The sequence was obtained with the Jalview software (Version 2.11.40).

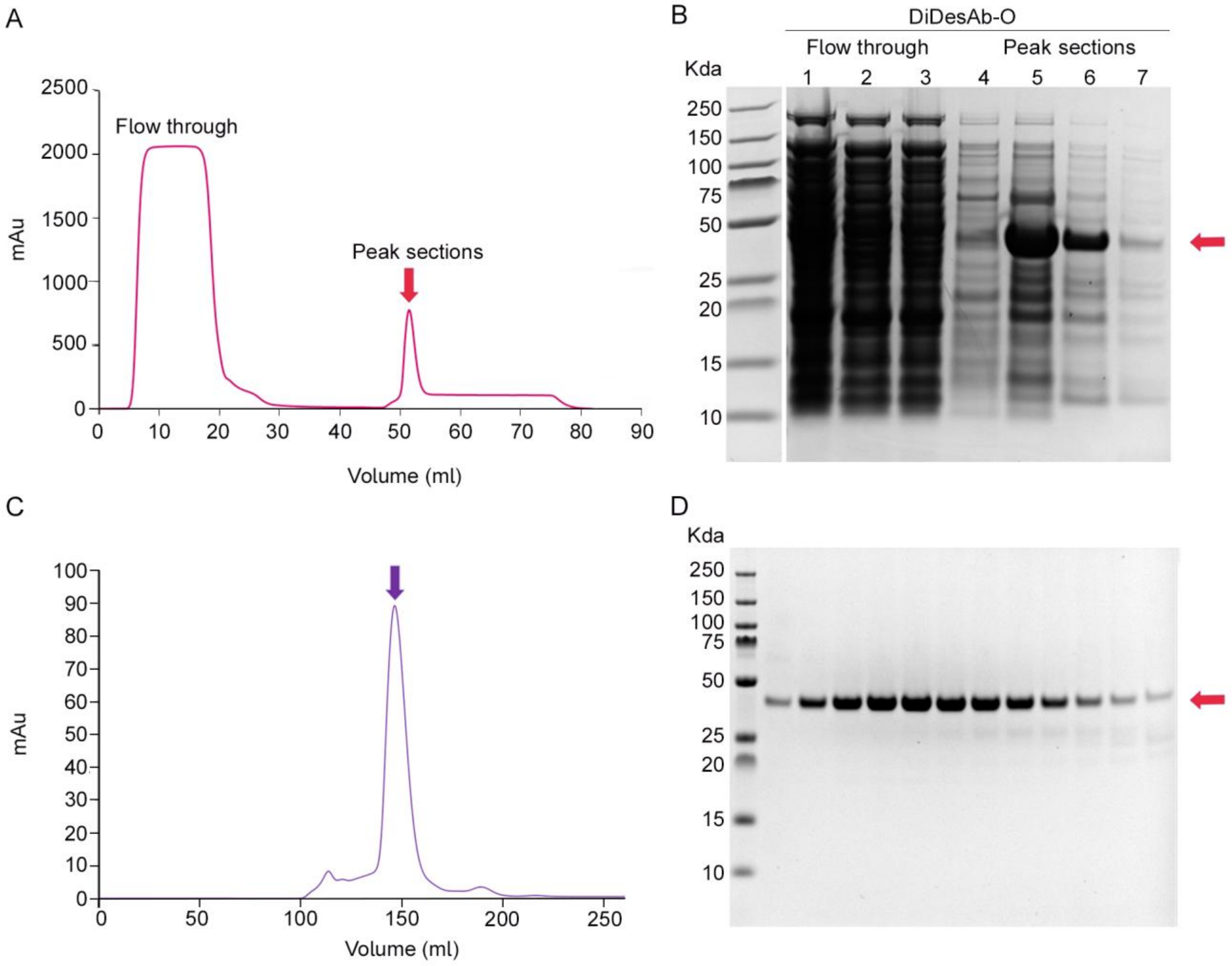


**Figure S2. DiDesAb-O expression and purification. A** $Ni^{2+}$-NTA chromatogram of lysed *E. coli* supernatant containing DiDesAb-O (red arrow). **B** SDS-PAGE of samples taken after $Ni^{2+}$-NTA chromatography; flow through (lanes 1-3), peak fractions (lane 4-7). DiDesAb-O molecular weight is indicated by the red arrow. **C** SEC chromatogram of the sample fractions after $Ni^{2+}$-NTA chromatography and containing DiDesAb-O. The peak relative to DiDesAb-O is indicated by the purple arrow. **D** SDS-PAGE of DiDesAb-O peak fractions taken after SEC to verify the sample purity. DiDesAb-O molecular weight is indicated by the red arrow.

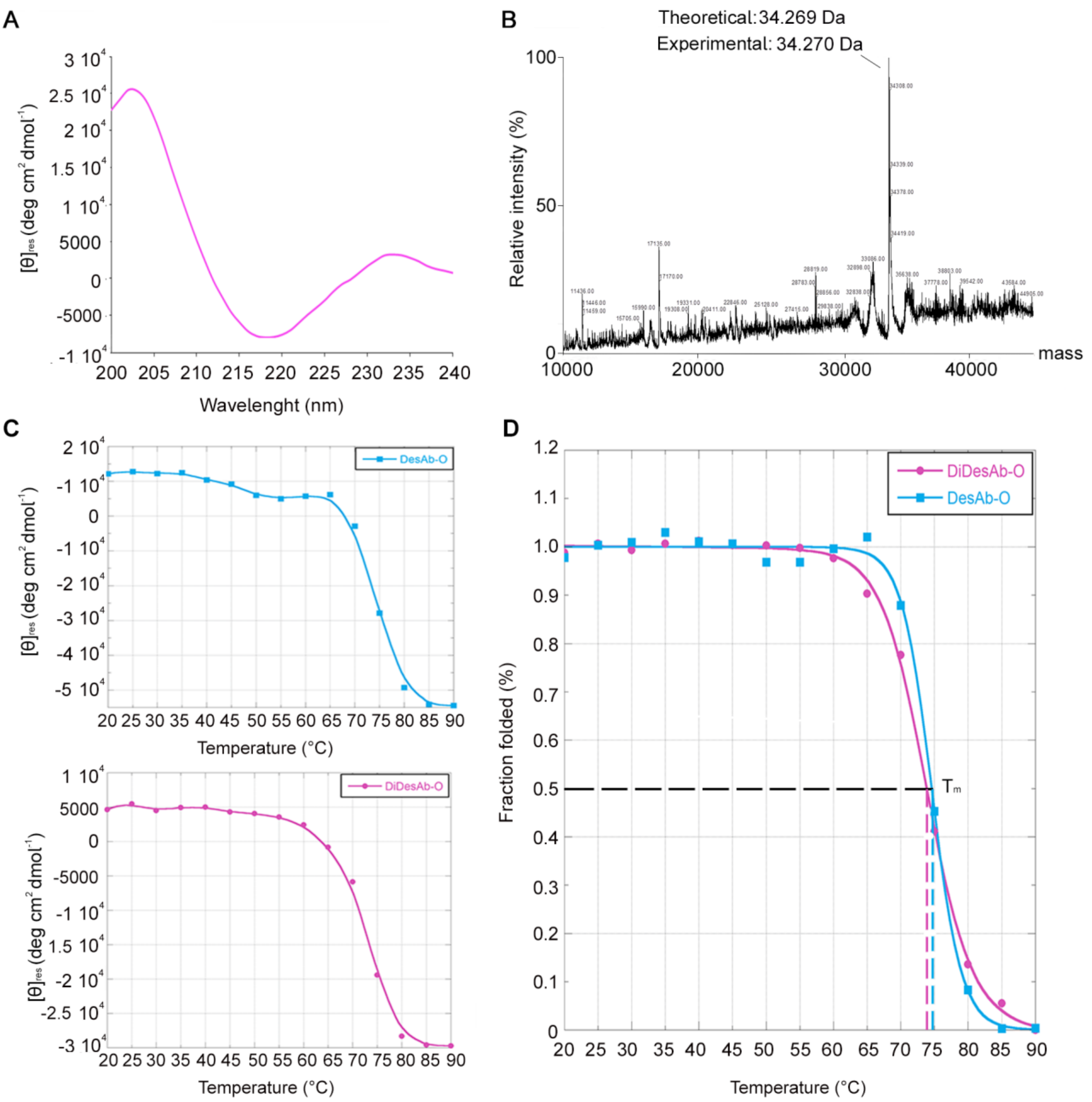


**Figure S3**. **Structural characterization of DiDesAb-O.** **A** Far-UV CD spectrum of DiDesAb-O. The presence of peaks at 202 nm and 218 nm highlights a predominant β-sheet secondary structure. **B** ESI-MS spectrum. Theoretical molecular weight was determined by Web tools such as ExPASy ProtParam tool (34.269 Da) being perfectly in line with the mass observed by ESI-MS (34.270 Da). **C** DesAb-O (top) and DiDesAb-O (bottom) denaturation curves were obtained using $[\theta]_{210}$ plotted against temperature between 20 °C and 90 °C. **D** DesAb-O and DiDesAb-O denaturation curves fitted with the Santoro and Bolen equation and normalized to fraction folded (%) values. DiDesAb-O had a temperature of half-denaturation ($T_m$) of 73.7 °C, while the value for DesAb-O was 74.6 °C.

| | DesAb-O (%) | DiDesAb-O (%) |
|---|---|---|
| **Helix 1 (regular)** | 0.0 | 0.0 |
| **Helix 2 (distorted)** | 0.0 | 2.6 |
| **Anti-parallel 1 (left-twisted)** | 6.4 | 8.8 |
| **Anti-parallel 2 (relaxed)** | 34.7 | 29.9 |
| **Anti-parallel 3 (right-twisted)** | 15.5 | 21.9 |
| **Parallel** | 6.9 | 0.0 |
| **Turn** | 8.8 | 10.4 |
| **Others** | 27.7 | 26.3 |

**Table S1.** Estimated secondary structure content (%) obtained with BestSel software (v1.3.230210; Micsonai et al., 2018) between the wavelength range of 195 and 250.

| $A\beta_{42}$:DiDesAb-O molar ratios | $t_{50}$ (h) |
|---|---|
| **1:1** | 3.2 ± 1.1 |
| **1:0.5** | 2.0 ± 0.6 |
| **1:0.25** | 1.8 ± 0.5 |
| **1:0.125** | 1.7 ± 0.5 |
| **1:0** | 1.5 ± 0.4 |

**Table S2**. $t_{50}$ (h) of solutions containing ThT and decreasing $A\beta_{42}$:DiDesAb-O molar ratios.

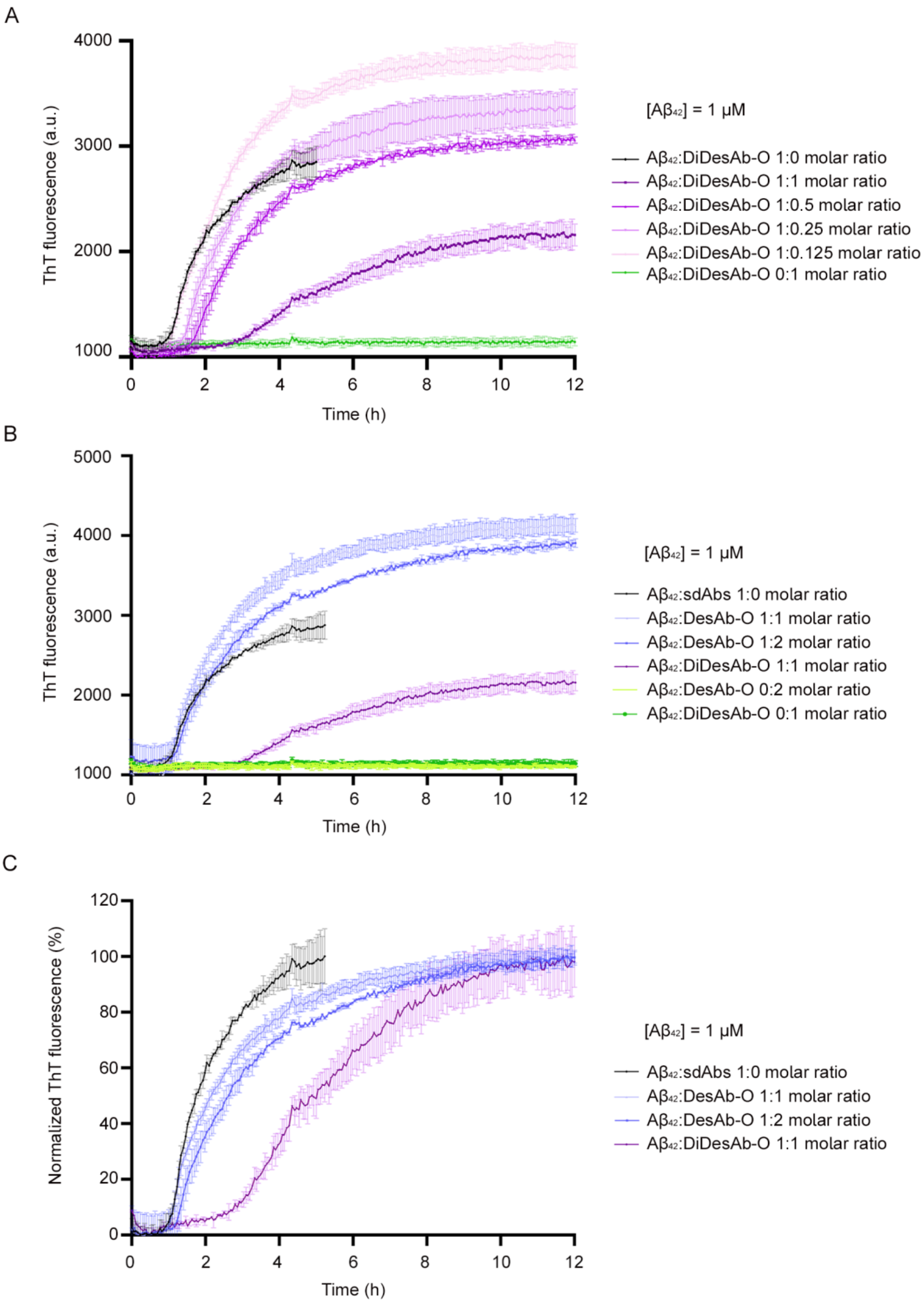


**Figure S4. Raw data of the ThT aggregation assays illustrated in Figure 3**. **A,B** Raw ThT aggregation data for 1 µM $A\beta_{42}$ aggregations at varying $A\beta_{42}$:DiDesAb-O **(A)** and $A\beta_{42}$:DesAb-O **(B)** ratios. Controls included DiDesAb-O and DesAb-O alone (1 µM and 2 µM, respectively). Two replicates were averaged for $A\beta_{42}$ alone, whereas three replicates were averaged for co-incubated solutions. **C** Normalization of the ThT assay kinetic traces represented in **B**. In all panels, error bars refer to standard deviations.

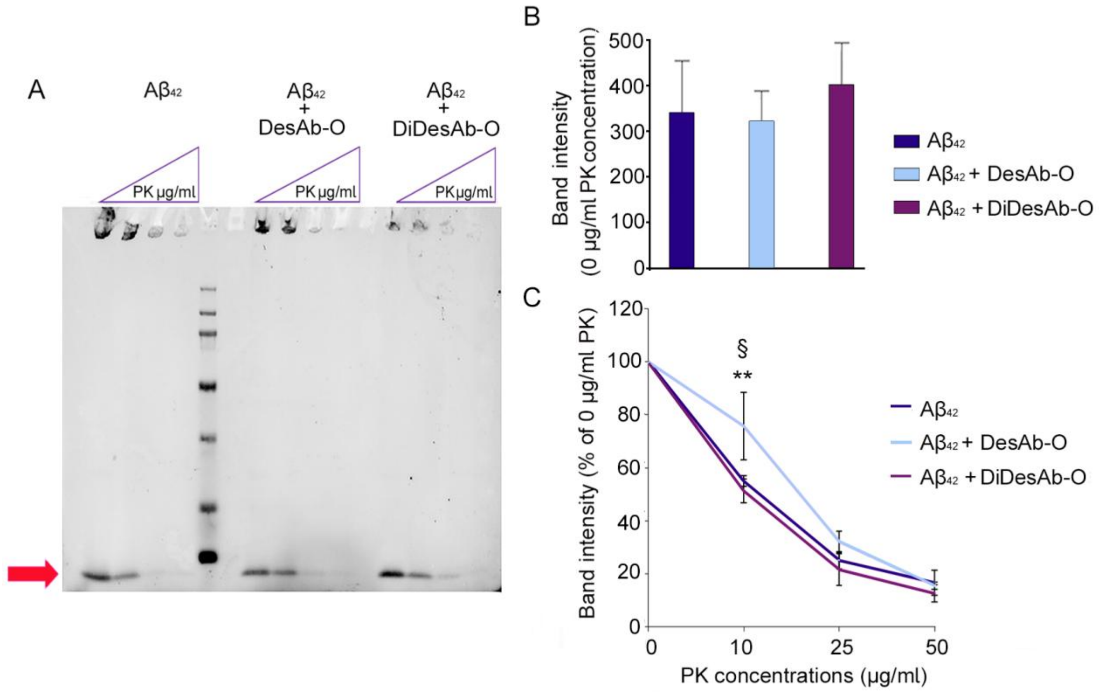


**Figure S5. Evaluation of PK cleavage sensitivity in A$\beta_{42}$ fibrils obtained in absence or presence of sdAbs. A** Representative Western Blot of A$\beta_{42}$ fibrils obtained after 4 days at 37 °C with or without co-incubation with sdAbs, treated with increasing PK concentrations (0, 10, 25, 50 $\mu$g/mL) for 30 mins. **B** Quantification of 0 $\mu$g/mL band intensity (n = 3) for each treatment. We can assess that all fibrils obtained in different conditions share similar band intensities at 0 $\mu$g/mL, demonstrating a consistent baseline across different treatment conditions, enabling reliable normalization of bands obtained at higher PK concentrations. **C** Evaluation of A$\beta_{42}$ fibril resistance to the PK digestion at increasing PK concentrations (0, 10, 25, 50 $\mu$g/mL). Experimental errors are S.E.M. Samples ($n = 3$) were analyzed by two-way ANOVA followed by Bonferroni's multiple-comparison test relative to Aβ fibrils obtained without sdAb versus Aβ fibrils obtained in the presence of DesAb-O (§$P < 0.05$), or to Aβ fibrils obtained in the presence of DiDesAb-O compared to Aβ aggregates obtained in the presence of DesAb-O (** $P < 0.01$).

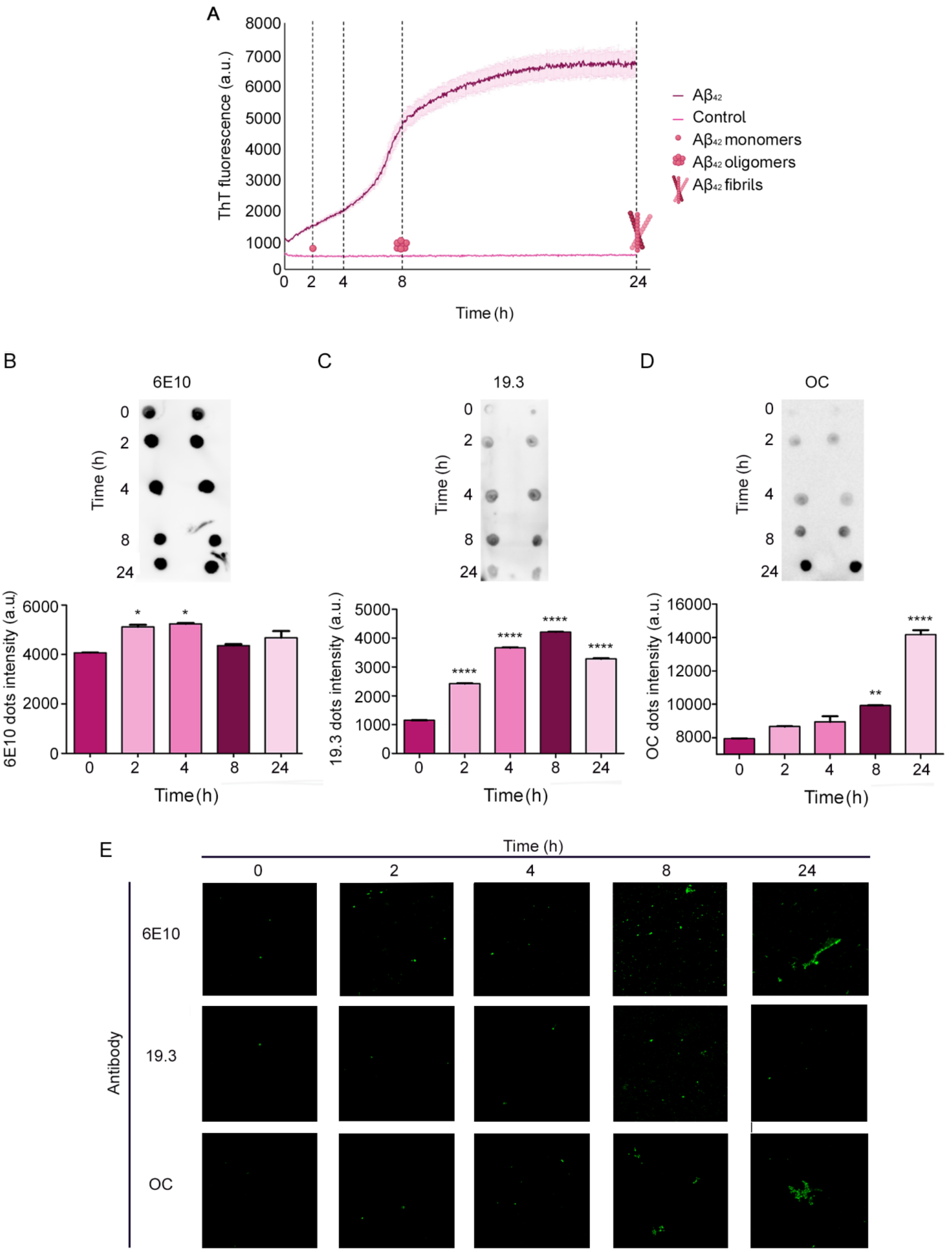


**Figure S6.** Characterisation of $A\beta_{42}$ aggregates starting from the synthetic peptide. **A** Monomeric $A\beta_{42}$ was incubated in PBS at 10 µM with 25 µM ThT dye, 37 °C. Time points were collected at 0, 2, 4, 8 and 24 h to perform a characterization of $A\beta_{42}$ aggregates. **B-D** Dot blot analysis and quantification of $A\beta_{42}$ samples collected at different timepoints. Samples of different $A\beta_{42}$ species were deposited (2 µl/spot) onto a nitrocellulose membrane and detected with the indicated antibodies (Abs). Membranes were incubated with 6E10 (**B**), 19.3 (**C**) and OC (**D**) primary Abs. Experimental errors are S.E.M. Samples (n = 2) were analysed by One-way ANOVA test followed by Bonferroni's multiple-comparison test relative to their respective time 0 h (*P < 0.05, **P < 0.01, and ****P < 0.0001). **E)** STED microscopy images and visualisation of Aβ$_{42}$ samples collected at different timepoints. Samples of different Aβ$_{42}$ species were immunolabeled with 6E10, 19.3 and OC Abs. The STED microscopy images are perfectly in line with the dot blot assay results, showing the presence of oligomers since the initial stage of the aggregation Aβ$_{42}$ process as represented in 0, 2 and 4 h of 19.3 Abs images. After 24 h, the presence of fibrils is proved by the OC Abs signal confirming the aggregation assay and dot blot assay results.

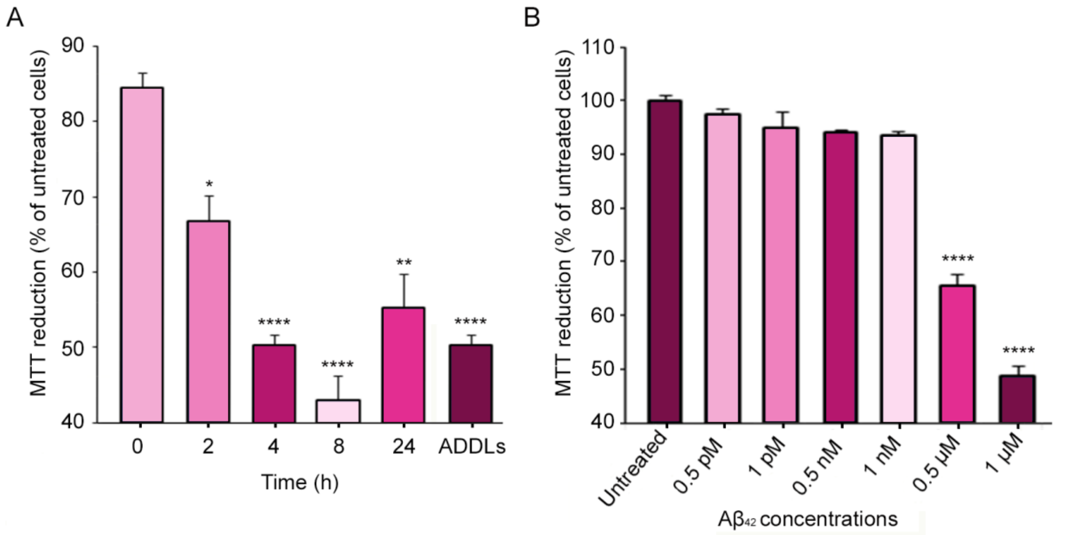


**Figure S7**. Aβ$_{42}$ oligomeric species obtained after 8 h of incubation exhibit high toxicity. **A** MTT reduction in SH-SY5Y cells treated for 24 h with various Aβ$_{42}$ aggregates (1 µM) collected at different timepoints (0, 2, 4, 8 and 24 h) during aggregation. ADDLs were used as a positive control. The most toxic aggregates were obtained after 8 h of aggregation, being in line with previous evidence of high Aβ$_{42}$ oligomers concentration at this timepoint (**Figure S6**). **B** MTT reduction in SH-SY5Y cells treated for 24 h with Aβ$_{42}$ aggregates obtained after 8 h of aggregation in a dose-dependent manner. Experimental errors are S.E.M. Samples (n = 2 and n = 3 for A and B, respectively) were analysed by One-way ANOVA followed by Bonferroni's multiple comparison test relative to untreated cells (*P<0.05, **P<0.01 and ****P<0.0001).
